# Evolution of regulatory networks associated with traits under selection in cichlids

**DOI:** 10.1101/496034

**Authors:** Tarang K. Mehta, Christopher Koch, Will Nash, Sara A. Knaack, Padhmanand Sudhakar, Marton Olbei, Sarah Bastkowski, Luca Penso-Dolfin, Tamas Korcsmaros, Wilfried Haerty, Sushmita Roy, Federica Di-Palma

## Abstract

Seminal studies of vertebrate protein evolution speculated that gene regulatory changes can drive anatomical innovations. However, very little is still known about gene regulatory network (GRN) evolution associated with phenotypic effect across ecologically-diverse species. Using a novel approach to reconstruct GRNs in vertebrate species, we aimed to study GRN evolution in representative species of the most striking example of an adaptive radiation, the East African cichlids. We previously demonstrated how the explosive phenotypic diversification of East African cichlids is attributed to diverse molecular mechanisms, including accelerated regulatory sequence evolution and gene expression divergence. To investigate these mechanisms across species at a genome-wide scale, our novel network-based approach identifies ancestral and extant gene co-expression modules along a phylogeny, and by integrating associated regulators, predicts candidate regulatory regions implicated in traits under selection in cichlids. As a case study, we present data from a well-studied adaptive trait - the visual system - for which we report striking cases of network rewiring for visual opsin genes, identify discrete regulatory variants, and investigate the plausibility of their association with cichlid visual system evolution. In regulatory regions of visual opsin genes, *in vitro* assays confirm that transcription factor binding site mutations disrupt regulatory edges across species, and segregate according to lake species phylogeny and ecology, suggesting GRN rewiring in radiating cichlids. Our approach revealed numerous novel potential candidate regulatory regions across cichlid genomes with no prior association, as well as those with previously reported associations to known adaptive evolutionary traits, thus providing proof of concept.

## Background

Seminal studies by King and Wilson[1] analysing protein evolution in vertebrates speculated the importance of evolutionary changes in ‘regulatory systems’ associated with morphological diversity[2,3]. These ideas were soon expanded on by François Jacob[4], who suggested that the molecular ‘tinkering’ of pre-existing systems is a hallmark of evolution where, for example, regulatory systems can either be transformed or combined for functional gain [4]. These theories underlie many studies on the divergence of regulatory systems associated with morphological evolution, and broadly focus on changes in gene regulatory networks (GRNs) that determine the expression patterns of genes [5,6]. Such changes can be mutations within transcription factor binding sites (TFBSs) located in *cis*-regulatory elements (promoters and enhancers) of genes or *trans* regulatory changes that are due to changes in the level of a regulator. Alterations of GRNs can ultimately lead to phenotypic divergence [7], and such changes between species are defined as ‘GRN rewiring events’. This is characterized by regulatory interactions present in one or more species but lost in another species, and potentially replaced by a new interaction between the orthologous TF and a target gene. Several comparative studies of GRNs underlying mechanisms of adaptation and evolution have been carried out in unicellular prokaryotes, *E. coli* [8] and several nonvertebrate eukaryotes, including yeast [9,10], plants [11], fruit fly [12] and echinoderms [12,13]. Whilst there are efforts to collate and integrate several genomic datasets for vertebrates, including human and mouse [14], reconstruction of regulatory networks from these data alone remains a major computational challenge and very little is known about the phenotypic effect of genome-wide regulatory network rewiring events in non-model vertebrates [15].

In vertebrates, ray-finned fishes are the largest radiation of any group, and amongst teleosts and even all vertebrates, the East African cichlid radiations represent arguably the most speciose modern example of an adaptive speciation. In the great lakes of East Africa (Victoria, Malawi and Tanganyika) and within the last few million years [16,17], one or a few ancestral lineages of haplochromine cichlid fish have given rise to over 1500 species. These species occupy a large diversity of ecological niches and differ dramatically in phenotypic traits, including skeletal morphology, dentition, colour patterning, and a range of behavioural traits. We have previously demonstrated that a number of molecular mechanisms have shaped East African cichlid genomes e.g. rapid evolution of regulatory elements, and the ‘evolutionary tinkering’ of these systems has provided the necessary substrate for diversification [18]. This, coupled with the recent origin of cichlid species and ongoing gene flow [19], suggests that evolutionary *cis*-regulatory changes have an important functional role in controlling gene expression and ultimately, phenotypic variation. Here we show how discrete *cis*-regulatory changes can fuel GRN rewiring in the evolution of cichlid radiations, and alongside several other molecular mechanisms, the ‘tinkering’ of regulatory systems could potentially serve as a substrate for the evolution of phenotypic diversity.

## Results

### Gene co-expression is tissue-specific and highlights functional evolutionary trajectories

We applied the Arboretum [9] algorithm to RNA-Seq data of six tissues in five species, and identified ten modules of 12,051-14,735 co-expressed genes (1,205-1,474 genes per module per species) (Fig. 1a). Modules of co-expressed genes across the five species show varying expression levels in specific tissues e.g. module 1 is eye specific, whilst module 3 is heart, kidney and muscle specific (Fig. 1a). Consistent with the phylogeny and divergence times, gene orthologs of the three closely related haplochromines (*P. nyererei, M. zebra* and *A. burtoni*) and *N. brichardi* are generally more conserved in module assignment than *O. niloticus* e.g. module 2, 4 and 6 (Fig. S-R1a, blue off-diagonal elements), indicating differential gene expression evolution across the species. Between the haplochromines, some gene orthologs are distributed in either one of two modules e.g. 0 or 8 (Fig. S-R1a, blue off-diagonal elements in haplochromines), indicating co-expression with different sets of genes and suggesting rewiring of transcriptional networks between cichlid species.

**Figure 1.**
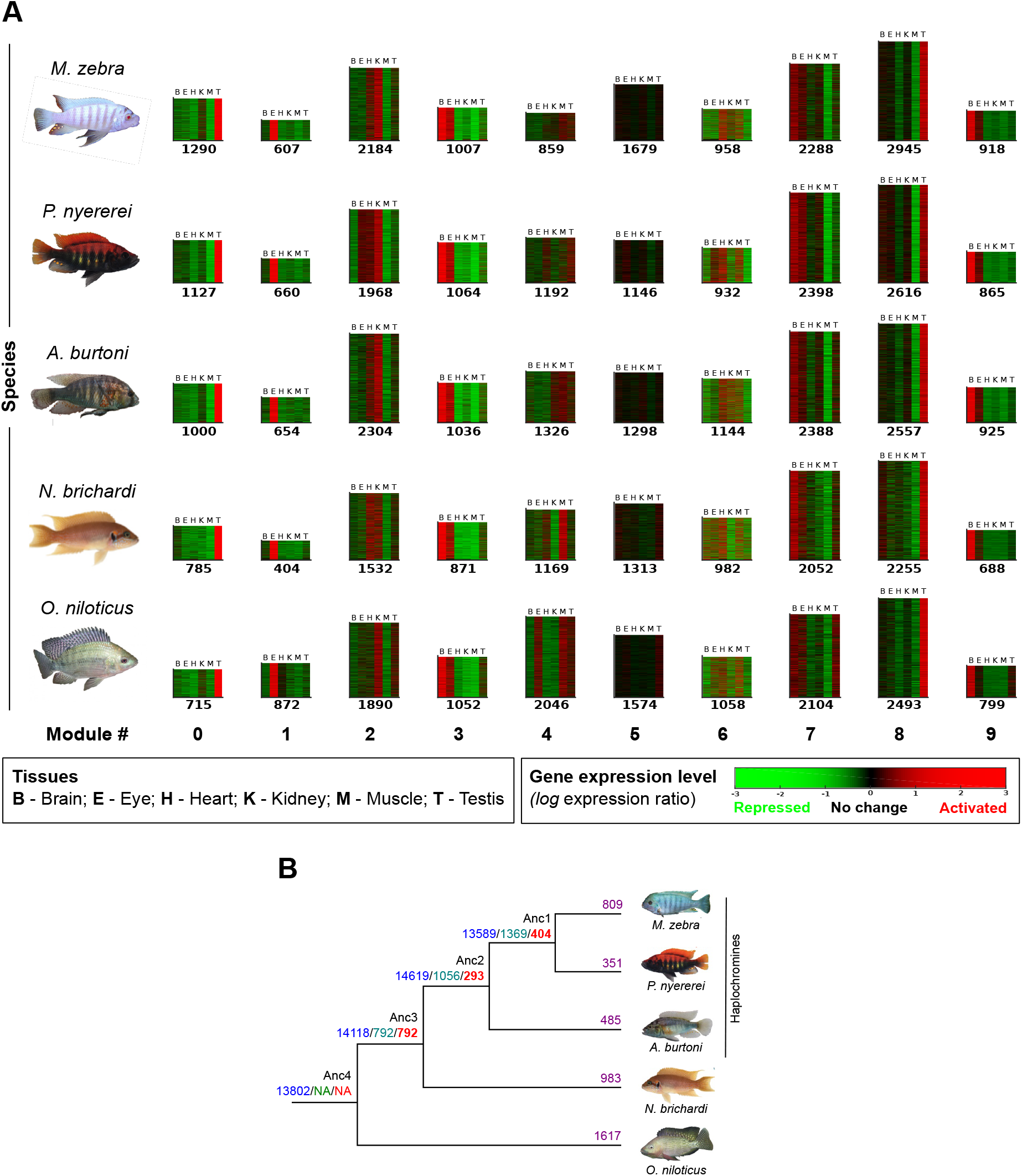
Evolution of gene expression in five cichlids. **(A)** 10 (0–9, heatmaps) coexpression modules identified by Arboretum [9] in six tissues of five cichlid species. Color bar denotes *log* expression ratio across each tissue, relative to the mean expression across all tissues - (red) activated; (green) repressed; and (black) no change. Each heatmap shows the expression profile of genes assigned to that module in a given species and height is proportional to number of genes in module (on *bottom*). **(B)** Number of state changes in module assignment along the five cichlid phylogeny [18]. Blue numbers: ancestral node genes assigned to modules; Green numbers: state changes compared to the deepest common ancestor (Anc4); Red numbers: state changes from last common ancestor (LCA); Purple numbers: state changes in the species compared to all other species.

In our approach, the assignment of co-expressed gene modules by Arboretum [9] is inferred using a probabilistic framework starting from the last common ancestor (LCA) in the phylogeny, allowing us to follow the evolution of gene co-expression along the species tree [9]. Orthologous genes of each species can be assigned to non-orthologous modules (Fig S-R1a), indicative of potential co-expression divergence and transcriptional rewiring from the LCA; referred to as ‘state changes’ in module assignment. We identified unique state changes and expression divergence of 655 genes along ancestral nodes (Fig. 1b), including several cellular and developmental TFs (51 TFs - Anc4/3; 20 TFs - Anc3/2; 34 TFs - Anc2/1) such as *foxo1, hoxa11* and *lbx1*. Several of these state changed regulatory TFs are also enriched (fold enrichment 1.1 - 1.7) in module gene promoters according to tissue-specific function; for example, promoters of module 1 genes (eye-specific expression) are significantly enriched (False Discovery Rate, FDR <0.05) for TF motifs involved in retina- and lens-related development/functions e.g. CRX, PITX3 and OTX1 [20] (Fig. S-R1c, Ext. Data S-R1B). Accordingly, we observe differences in the levels of TF motif enrichment across species genes, including retina/lens related TFs e.g. *RARα/β/γ* and *RXRα/β/γ* [21] of module 1 gene promoters in all species except *N. brichardi* (Fig. S-R1c, Ext. Data S-R1B). Differences in motif enrichment could be associated with TF expression changes, where state-changes (Fig. 1b) reflect shifted domains of tissue expression, and imply differential regulatory control of target genes across tissues and along the phylogeny. We test this by calculating the Pearson correlation coefficient (PCC) between cross-species TF motif enrichment and tissue expression for all 337 expressed TFs that have 2064 motif enrichments across module genes (Extended Data S-R1C-H). We note different patterns of correlation between cross-species TF motif enrichment and tissue-specific expression; 120-148 TFs had no correlation (PCC 0-0.01); including when there are large shifts in motif enrichment and/or expression in several species (several phylogenetic state-changes) e.g. Kidney-Module2-FOXO1 (PCC=0.01) (Extended Data S-R1F). On the other hand, there is positive correlation ranging from small (PCC 0.1-0.3) for 225-271 TFs, medium (PCC 0.3-0.5) for 211-274 TFs, and large (PCC 0.5-1) for 351-437 TFs; this includes cases where there is comparable motif enrichment across species and either no shifts (no TF state-changes) e.g. Brain-Module9-FOXA2 (PCC=0.97) or focused shifts (TF state-change in one or subsets of species) e.g. Eye-Module2-CDX1 (PCC=0.98) in TF tissue expression (Fig. S-R1e, Extended Data S-R1C-H). Focusing on modules associated with specific tissues, e.g. module 1 which contains eye-expressed genes, we find that retinal TFs that are known to modulate opsin expression e.g. CRX [22], have variable motif enrichment (fold enrichment 1.2 – 1.4) in eye-expressed genes, that is associated (PCC=0.85) with a concurrent change (increase or decrease) in TF eye expression along the phylogeny (Fig. S-R1f; see *Supplementary Text*). For most TFs (226-262/337 TFs) and tissues, motif enrichment is largely correlated (PCC 0.5 - 1) with TF expression; similar motif enrichment (across species) is associated with either expression conservation (across all species) or subtle expression changes (in one or subsets of species), and is more stable (in expression differences) than TFs with variable motif enrichment along the phylogeny (Extended Data S-R1C-H). Overall, this highlights that gene co-expression differences between species could be driven by differences in gene regulatory programmes, and could be associated with regulatory network changes underpinning traits under selection in cichlids, such as the visual system [23].

### Fine scale nucleotide variation at TF binding sites drives regulatory divergence in cichlids through GRN rewiring

Central to gene expression regulation are *cis*-regulatory elements (including promoters and enhancers) that harbour several transcription factor binding sites (TFBSs) and mutations thereof, can drive GRN evolution, altering target gene transcription without affecting the expression pattern of genes co-regulated by the same TF. In the five cichlid genomes, there is no significant increase in evolutionary rate at promoter regions compared to fourfold-degenerate sites (Fig. S-R2a) however, we identify a few outlier genes with significantly higher evolutionary rate at promoter regions at ancestral nodes (12-351 genes, Fig. S-R2aB) and within species (29-352 genes, Fig. S-R2aD), indicative of small-scale changes in promoter regions (see *Supplementary Text*). Of all the identified pairwise species variation (8 to 32 million variants), a significant proportion (13-28%) overlap predicted TFBSs in promoter regions, and this is higher than (8-9%) of variants overlapping neutrally evolving regions flanking gene promoters (Table S-R2a, Fig. S-R2b). GO enrichment analysis of co-expressed genes with variation in their regulatory regions, against a background of all genes in each genome, highlight associations with key molecular processes e.g. signal transduction - non-state changed promoter TFBSs (Fig. S-R2c). These findings imply that differences in gene regulatory programmes are likely driven by discrete nucleotide variation at regulatory binding sites, and could have functional implications in cichlids.

To further investigate patterns of divergent regulatory programmes that could be associated with the observed changes in regulatory binding sites, we developed and applied a computational framework (see *Methods*, Fig. S-M1) to study regulatory relationships; this involved the reconstruction of species-specific GRNs through the integration of different genomic datasets (Table S-R3a). We focused on regulatory relationships with DNA; this involved integrating an expression-based network with *in-silico* predictions of TF binding to target gene (TG) promoters using our cichlid-specific and vertebrate-wide TF motif scanning pipeline (see *Methods*, Fig. S-M1). We first used species- and module-specific gene expression levels to infer an expression-based network [24] (see *Methods*, Fig. S-M1) of 3,180-4,099 transcription factor - target gene (TF-TG) edges across the five species (FDR<0.05, Table S-R3a). Next, based on our *in-silico* TFBS motif prediction pipeline, we predicted TFBS motifs up to 20kb upstream of a genes transcription start site (TSS), and using sliding window analysis of 100 nucleotides (nt), we retained TF motifs in the gene promoter region, defined as up to 5kb upstream of a genes TSS (see *Methods*, Fig. S-M2). Each statistically significant TFBS motif (FDR<0.05) was associated to its proximal target gene (TG), and represented as two nodes and one TF-TG edge. Based on the integrated approach (see *Methods*, Fig. S-M1), we predicted a total of 3,295,212-5,900,174 TF-TG edges (FDR<0.05) across the five species, that could be encoded into a matrix of 1,131,812 predicted TF-TG edges (FDR<0.05) present in at least one species. To ensure accurate analysis of GRN rewiring through an integrative approach, all collated edges were then further pruned to a total of 843,168 TF-TG edges (FDR<0.05) where the edge is 1) present in at least one species; 2) not absent in at least one species due to gene loss/poor annotation; and 3) based on the presence of nodes in modules of co-expression genes (see *Methods*).

We used three metrics to study large-scale TF-TG network rewiring between species, that included: 1) state changes in module assignment; 2) DyNet [25] network rewiring scores; and 3) TF rate of edge gain and loss in networks. The first metric compares TF-TG edges of a “focal” species versus the other species in the context of gene coexpression, while the second and third metric compute a likelihood score for the overall extent (across all species) of edge changes associated with nodes of a single gene of interest. We first focused on 6,844 1-to-1 orthologous genes represented in 215,810 TF-TG interactions, termed ‘TF-TG 1-to-1 edges’, along the five cichlid tree. Using a background set of all module genes (18,799 orthogroups), the TF-TG 1-to-1 edges are associated with morphogenesis and cichlid traits under selection e.g. eye and brain development (FDR <0.05, Fig. S-R3aA). There are 379 TFs represented in the TF-TG 1-to-1 edges, and we focus on their interactions to determine whether TFs with (state) changes in module assignment show altered regulatory interactions (rewiring in a focal vs other species). Of the 379 TFs, 13-18% (50-70 TFs) are rewired (spanning 4,060-9,423/215,810 edges, FDR<0.05, Fig. 2a; see *Supplementary Text*) and change module assignment across the five species (in 1 focal vs all 4 other species). Their altered TF-TG edges are associated with signalling pathways and processes such as cell differentiation and embryonic development (FDR<0.05, background of all module genes, Fig. 2b). Further examination of rewiring rates in the networks of 6,844 1-to-1 orthologous genes (in 215,810 TF-TG interactions) using the DyNet [25] degree-corrected rewiring (*D_n_*) score (Fig. 2c, Extended Data Table S-R3A) identifies 9 teleost and cichlid trait genes associated with morphogenesis from previous studies (Fig. 2c, Extended Data Table S-R3B). These genes have a few standard deviations higher degree-corrected rewiring (*D_n_*) score than the mean (0.17±0.03 SD) of all 1-to-1 orthologs and their rewiring score is higher (Kolmogorov–Smirnov KS-test *p-value* = 6 × 10^−4^) (Fig. 2c – left violin plot, orange dots; Extended Data Table S-R3C; see *Supplementary Text*). Examples of those genes include *gdf10b* associated with axonal outgrowth and fast evolving in cichlids [18] and the visual opsin gene, *rh2* (Fig. 2c – left violin plot; Extended Data Table S-R3C). We extend our analyses beyond the 6,844 1-to-1 orthologs only, by including an additional 7,746 many-to-many orthogroups (see *Methods*) resulting in a set of 843,168 ‘TF-TG all edges’ across the five species. Using a background set of all module genes (18,799 orthogroups), the genes in these TF-TG edges are associated with morphogenesis e.g. retina development (FDR<0.05, Fig. SR3aB). These edges include interactions of 783 TFs of which, 13-18% (100-140 TFs) are predicted to be rewired (in 20,716-37,590/843,168 edges, FDR<0.05, Fig. 2d) and change module assignment across the five species (in 1 focal vs all 4 other species), indicating their associated transcriptional programs (FDR<0.05, background of all module genes) are also altered (Fig. 2e). By examining the network rewiring rates of 14,590 orthogroups (in 843,168 TF-TG interactions, Extended Data Table S-R3D), we identify 60 teleost and cichlid trait genes associated with phenotypic diversity from previous studies (Fig. 2c – *right* violin plot; Extended Data Table S-R3E). These genes have a few standard deviations higher degree-corrected rewiring (*D_n_*) score than the mean (0.23±0.007 SD) of all orthologs, and their rewiring score is higher (KS-test *p-value* = 6 × 10^−14^) (Fig. 2c – *right* violin plot, orange dots; Extended Data Table S-R3D). These genes include those associated with craniofacial development e.g. *dlx1a* and *nkx2-5* [20], telencephalon diversity e.g. *foxg1* [26], tooth morphogenesis e.g. *notch1* [27] and strikingly, most visual opsins e.g. *rho, sws2* and *sws1*, as well as genes associated with photoreceptor cell differentiation, *actr1b* [28] and eye development, *pax6a* [20] (Fig. 2c – *right* violin plot; Extended Data Table S-R3E). We then focus on the gain and loss rates of 186/783 TFs with >25 TF-TG edges along the five cichlid tree (see *Methods*). Out of the 186 TFs, 133 (72%) are predicted to have a higher rate of edge gain than loss e.g. DLX5 and NEUROD2, possibly acting as recruited regulators of gene expression in each branch from their last common ancestor (LCA) (Extended Data Table S-R3F); whereas 53/186 TFs (28%) have a higher loss of edges than gains e.g. OLIG2 and NR2C2, implying loss of gene expression regulatory activity from their LCA (Extended Data Table S-R3F). This suggests that along the five cichlid tree, TFs and their binding sites are evolving towards gaining, rather than losing regulatory edges, and possibly regulatory activity of genes from their LCA.

**Figure 2 –.**
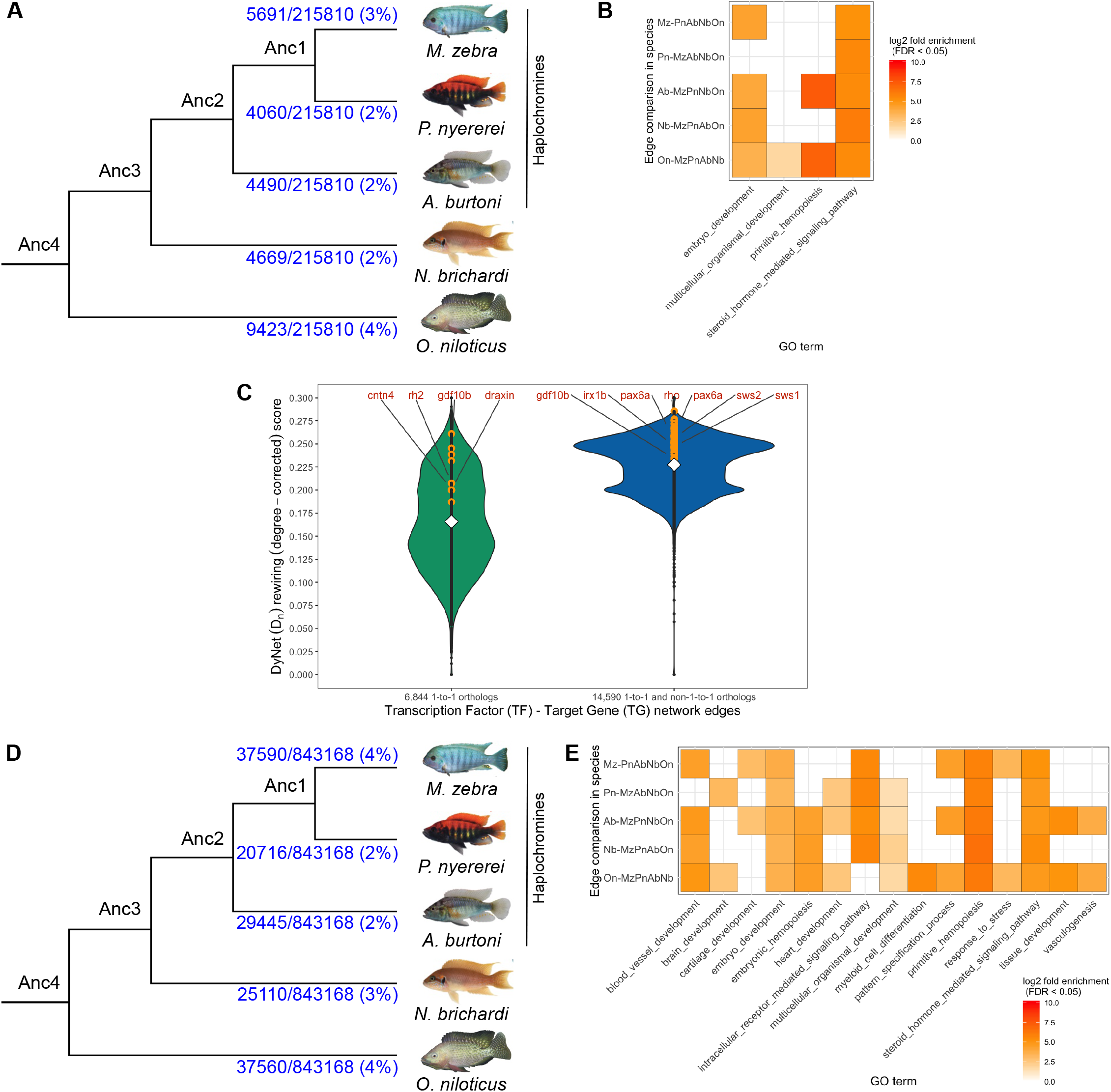
Network rewiring between species including TFs state changing coexpression module assignment and their targets (TGs). **(A)** Re-wiring events linked to module assignment state changes of TFs in 215,810 TF-TG 1-to-1 edges (FDR<0.05) of each species in the cichlid phylogeny compared to the other four species (see *Supplementary Text* for other FDR thresholds). **(B)** GO term enrichment of 50-70 rewired and state changed TFs and their associated TGs in each focal species vs the other 4 species (4,060-9,423/215,810 TF-TG 1-to-1 edges, Fig. 2A) against a background of all module genes, shown as grid heatmap of *log10* fold enrichment (legend on *right*, FDR <0.05). **(C)** Violin plots of DyNet (Dn) rewiring score (degree-corrected) from 6,844 1-to-1 orthologs in 215,810 TF-TG network edges (green, left violin) and 14,590 1-to-1 and many-to-many orthologs in 843,168 TF-TG network edges (blue, right violin). Mean rewiring score shown within each plot (white diamond). Degree-corrected rewiring score shown for non-candidate genes (black dots through center) and candidate morphogenetic trait genes (orange dots) with rewiring scores higher than the mean, and selected candidate examples are demarcated within. **(D)** Re-wiring events linked to module assignment state changes of TFs in 843,168 TF-TG all edges of each species in the cichlid phylogeny against the other four species. **(E)** GO term enrichment of 100-140 re-wired and state changed TFs and their associated TGs in each focal species vs the other 4 species (20,716-37,590/843,168 edges TF-TG all edges, Fig. 2D) against a background of all module genes, shown as grid heatmap of *log10* fold enrichment (legend on *right*, FDR <0.05).

To further characterise the role of the observed changes in *cis*-regulatory elements and their potential association with cichlid traits, we extended our analyses to include several radiating cichlid species. We screened all predicted TFBS (see *Methods*) variants between *M. zebra* and the other four cichlids, with their corresponding positions in 73 phenotypically distinct Lake Malawi species [19], to identify between species variation at regulatory sites along the phylogeny (Fig. S-R3b). As expected, the majority of variation at regulatory sites is identified between *M. zebra* and distantly-related Lake Malawi species clades e.g. NKX2.1 TFBS in *sws1* gene promoter, whereas shared ancestral sites are found with mainly same/closely-related Lake Malawi clades e.g. EGR2 TFBS in *cntn4* gene promoter. This indicates that genes associated with traits under selection e.g. visual systems [23] (*sws1*) and morphogenesis [18] (*cntn4*), harbour between species regulatory variants that segregate according to phylogeny and ecology of radiating lake species.

### *Cis*-regulatory changes lead to GRN alterations that segregate according to phylogeny and ecology of radiating cichlids

Through our integrative approach, we can examine the regulatory network topology of several genes associated with unique cichlid traits [29,30] represented by our six tissues. As a case study, we focus on the cichlid visual system; the evolution of cichlid GRNs and diverse palettes of co-expressed opsins can induce large shifts in adaptive spectral sensitivity of adult cichlids [23] and thus, we hypothesise that opsin expression diversity is the result of rapid adaptive GRN evolution in cichlids. Indeed, by focusing on species utilizing the same wavelength visual palette and opsin genes, we previously noted that several visual opsin genes (*rh2b, sws1, sws2a* and *rho*) have highly rewired regulatory networks (Extended Data Table S-R3F). Across the predicted transcriptional networks of cichlid visual opsins, there are several visual-system associated TF regulators of opsin genes (*sws2a, rh2b* and *rho*) that are either common e.g. STAT1A, CRX and GATA2, or unique to each species e.g. IRF1, MAFA and GATA2A (Fig. S-R4a-c). Such patterns of TF regulatory divergence could contribute to differential opsin expression, important for peak absorption spectra of the regulated opsin and ecology of each species.

*Sws1* (ultraviolet) opsin is utilized as part of the short-wavelength sensitive palette in *N. brichardi* and *M. zebra*. Whilst there are common regulators associated with retinal ganglion cell patterning in both species networks e.g. SATB1 [31], as well as several unique regulators associated with nuclear receptor signalling e.g. RXRB and NR2C2 [32] and retinal neuron synaptic activity e.g. ATRX [33] (Fig. 3a), the *sws1* networks of these two species have substantially diverged. Overall, using an FDR<0.05 for predicted edges, there are more predicted unique TF regulators of *sws1* in *M. zebra* (38 TFs) as compared to *N. brichardi* (6 TFs) (Fig. 3a, *bottom right*). Such tight TF-based regulation of *N. brichardi sws1* could induce rapid shifts in expression and spectral shift sensitivities between a larger peak ƛ_max_ of 417 nm in *N. brichardi* single cones [34] compared to 368 nm of *M. zebra* Sws1 [35]. Also, diverse regulation in *M. zebra* can increase *sws1* expression and in turn, increase spectral sensitivity to UV light and the ability for *M. zebra* to detect/feed on UV-absorbing phytoplankton and algae, as previously shown for Lake Malawi cichlids [36]. Furthermore, we identify that a candidate regulatory variant has likely broken the *M. zebra* NR2C2/RXRB shared motif that is otherwise predicted

**Figure 3.**
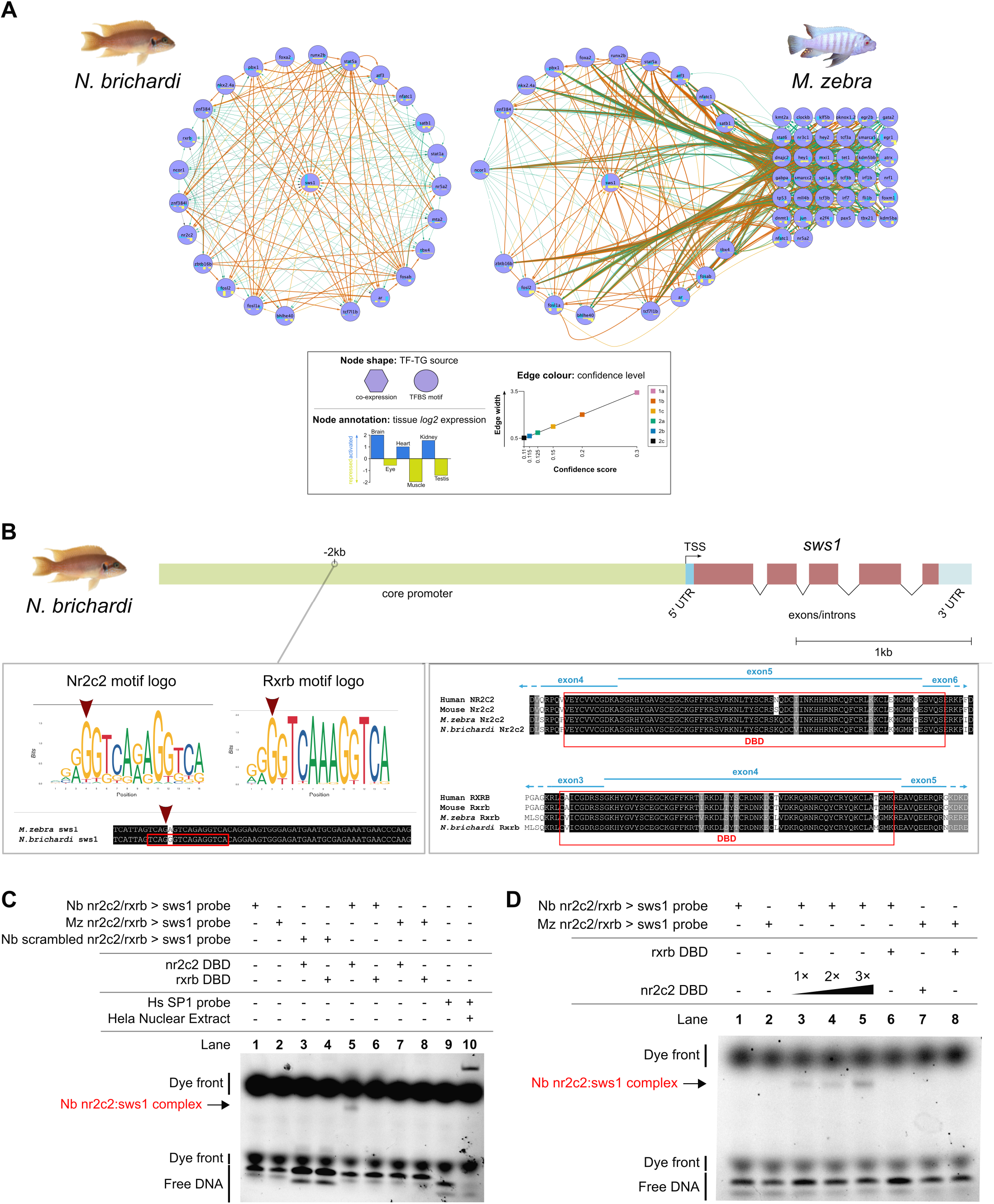
Evolution of the *sws1* opsin regulatory networks in *N. brichardi* and *M. zebra*. **(A)** Reconstructed regulatory networks of *sws1* opsin shown for *N. brichardi (left*) and *M. zebra* (*right*): circular layout nodes are common regulators (unless missing); grid layout nodes are unique regulators. Node shape, annotation and edge color denoted in legend to *left bottom*. Violin plot of significance (FDR<0.05) of unique TF-*sws1* edges in *N. brichardi (green violin*) and *M. zebra* (*blue violin*) to *bottom right* - mean edge significance score shown within each plot (white diamond); edges more than the mean (less significant) are shown as grey dots, and edges less than the mean (more significant) are shown as orange dots; selected example TFs are demarcated within. **(B)** On *left*, NR2C2 and RXRB motif logos and motif prediction in negative orientation *N. brichardi sws1* gene promoter (red box) and variant in *M. zebra sws1* gene promoter (red arrow). On *right*, NR2C2 and RXRB partial protein alignment showing DNA-binding domain (DBD) annotation in human, mouse, *M. zebra* and *N. brichardi*. **(C)** EMSA validation of NR2C2 and RXRB DBD binding to *N. brichardi* and *M. zebra sws1* gene promoter. Table denotes combinations of DNA probe and expressed DBD in EMSA reactions that include negative controls (lane 1 to 4); *N. brichardi* binding positive control (lane 5 and 6); *M. zebra* binding positive control (lane 7 and 8); kit negative (lane 9) and binding positive control (lane 10). **(D)** EMSA validation of increasing Nr2c2 DBD concentrations and binding to predicted TFBS in *N. brichardi sws1* gene promoter.

2kb upstream of the *N. brichardi sws1* TSS (Fig. 3b). Functional validation via EMSA confirms that NR2C2 and not RXRB binds to the predicted motif in the *N. brichardi sws1* promoter, forming a complex, and the variant has likely disrupted binding, and possibly regulation of *M. zebra sws1* (Fig. 3c-d). This is further supported by correlating expression values of these regulators and *sws1*, where NR2C2 is better associated with *sws1* than RXRB, particularly when focusing on eye tissue (Fig. S-R4eA *on right;* Fig. S-R4eB; see *Supplementary text*).

In another example, *rhodopsin (rho*), associated with dim-light vision is predicted to be regulated by GATA2 in *O. niloticus, A. burtoni* and *M. zebra* but not its duplicate gene, GATA2A only in *M. zebra* (Fig. S-R4c). We identify a candidate variant (red arrow, Fig. 4a) that has likely broken the *M. zebra* GATA2A motif that is otherwise predicted 1.8kb and 1.9kb upstream of the *O. niloticus* and *A. burtoni rho* TSS (Fig. 4a). Functional validation via EMSA confirms that GATA2A binds to the predicted motif in the *O. niloticus* and *A. burtoni rho* promoter, and the variant is likely to have disrupted binding, and possibly regulation of *M. zebra rho* (Fig. 4b). Species-specific expression correlations with the *rho* target gene are supportive of GATA2’s possible conserved role in all three species (*O. niloticus* PCC=0.89; *A. burtoni* PCC=0.39; *M. zebra* PCC=0.28; Fig. S-R4fC). While a more divergent role of GATA2A (O. niloticus PCC=0.79 and A. burtoni PCC=0.21) and negative correlation in *M. zebra* (PCC=-0.18) is supportive (Fig. S-R4fC) of the EMSA validation (Fig. 4). Based on these results and the known effects of regulatory mutations on cichlid opsin expression [37,38], discrete point mutations in TFBSs could be driving GRN evolution and rewiring events in traits that are under selection in radiating cichlids.

**Figure 4.**
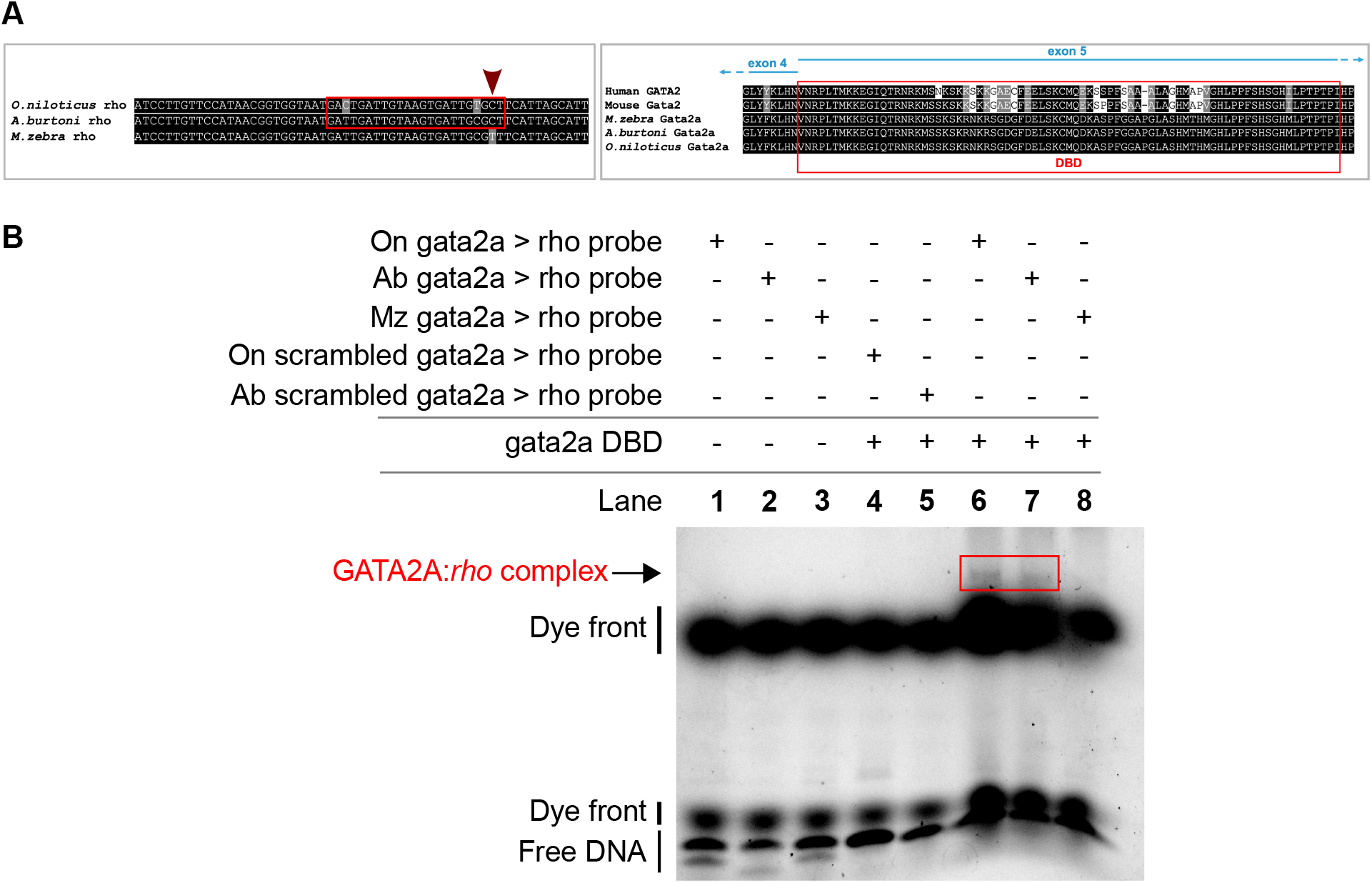
Evolution of the *rhodopsin* regulatory networks in *O. niloticus, A. burtoni* and *M. zebra*. **(A)** On *left*, GATA2A motif prediction in reverse orientation *O. niloticus* and *A. burtoni rhodopsin* gene promoter (red box) and substitution in *M. zebra rhodopsin* gene promoter (red arrow). On *right*, GATA2A partial protein alignment showing DNA-binding domain (DBD) annotation in human, mouse, *O. niloticus, A. burtoni* and *M. zebra*. **(B)** EMSA validation of GATA2A DBD binding to *O. niloticus* and *A. burtoni rhodopsin* gene promoter. Table denotes combinations of DNA probe and expressed DBD in EMSA reactions that include negative controls (lane 1 to 5); *O. niloticus* (lane 6), *A. burtoni* (lane 7) and *M. zebra* (lane 8) binding positive controls. GATA2A:*rho* complex formed in *O. niloticus* (lane 6) and *A. burtoni* (lane 7) as confirmed by band shift (red box) and no complex formed in *M. zebra* (lane 8).

Finally, we studied GRN rewiring as a result of between species TFBS variation in the context of phylogeny and ecology of lake species. Owing to the variability and importance of spectral tuning of visual systems to the foraging habits of all cichlid species, we focused on variants at regulatory sites of rewired visual opsin genes in the Lake Malawi species, *M. zebra*, as a reference to compare GRN rewiring (through TFBS variation) that could be associated with the ecology of sequenced Lake Malawi species [19]. If indeed the TFBSs are likely functional, we hypothesise that radiating species with similar foraging habits would share conserved regulatory genotypes, to possibly regulate and tune similar spectral sensitivities; whereas distally related species with dissimilar foraging habits would segregate at the corresponding regulatory site. Using 157,232 sites with 1) identified variation between the five cichlid species; and 2) are located in TFBSs of *M. zebra* candidate gene promoters; we identified 5,710 of the sites with between species variation across 73 Lake Malawi species (Fig. S-R3b), and flanking sequence conservation, representative of shared ancestral variation. The homozygous variant (T|T) that breaks binding of NR2C2 to *M. zebra sws1* promoter (Fig. 3, Fig. 5 *blue arrow*) is 1) conserved with the fellow algae eater, *T. tropheops* that also utilizes the same short-wavelength palette; 2) heterozygous segregating (*P. genalutea* - C|T and *I. sprengerae* - T|C) in closely related Mbuna species; and 3) homozygous segregated (C|C) in distantly related Mbuna species (*C. afra, C. axelrodi* and *G. mento*) and most other Lake Malawi species of which, some utilize the same short-wavelength palette and are algae eaters e.g. *H. oxyrhynchus* (Fig. 5). This suggests that in species closely related to *M. zebra*, and with a similar diet and more importantly, habitat, *sws1* may not be regulated by NR2C2, whilst in other species it could be, similar to *N. brichardi* (Fig. 3, Fig. 5 *red arrow*). In another example, regulation of *rho* by GATA2, and not its duplicate, GATA2A (Fig. 4), could be sufficient for regulating dim-light vision response in some rock dweller species (*M. zebra* and possibly *P. genulatea, T. tropheops* and *I. sprengerae*) but both *gata2* copies could be required to regulate *rho* in many other Lake Malawi species (79% with C|C genotype that otherwise predicts the GATA2A TFBS in *rho* gene promoter), as well as *A. burtoni* and *O. niloticus* (Fig. S-R4c, d). This highlights the potential differential usage of a duplicate TF in dim-light vision regulation. Based on these examples of TFBS variants that segregate according to phylogeny and ecology of lake species, GRN rewiring through TFBS variation could be a key contributing mechanism of evolutionary innovation in East African cichlid radiations. Such differences in visual tuning could correspond to species variation of habitat choice, foraging habits, diet, and male nuptial colouration. However, this requires further experimental testing, particularly in phenotypically divergent species pairs.

**Figure 5.**
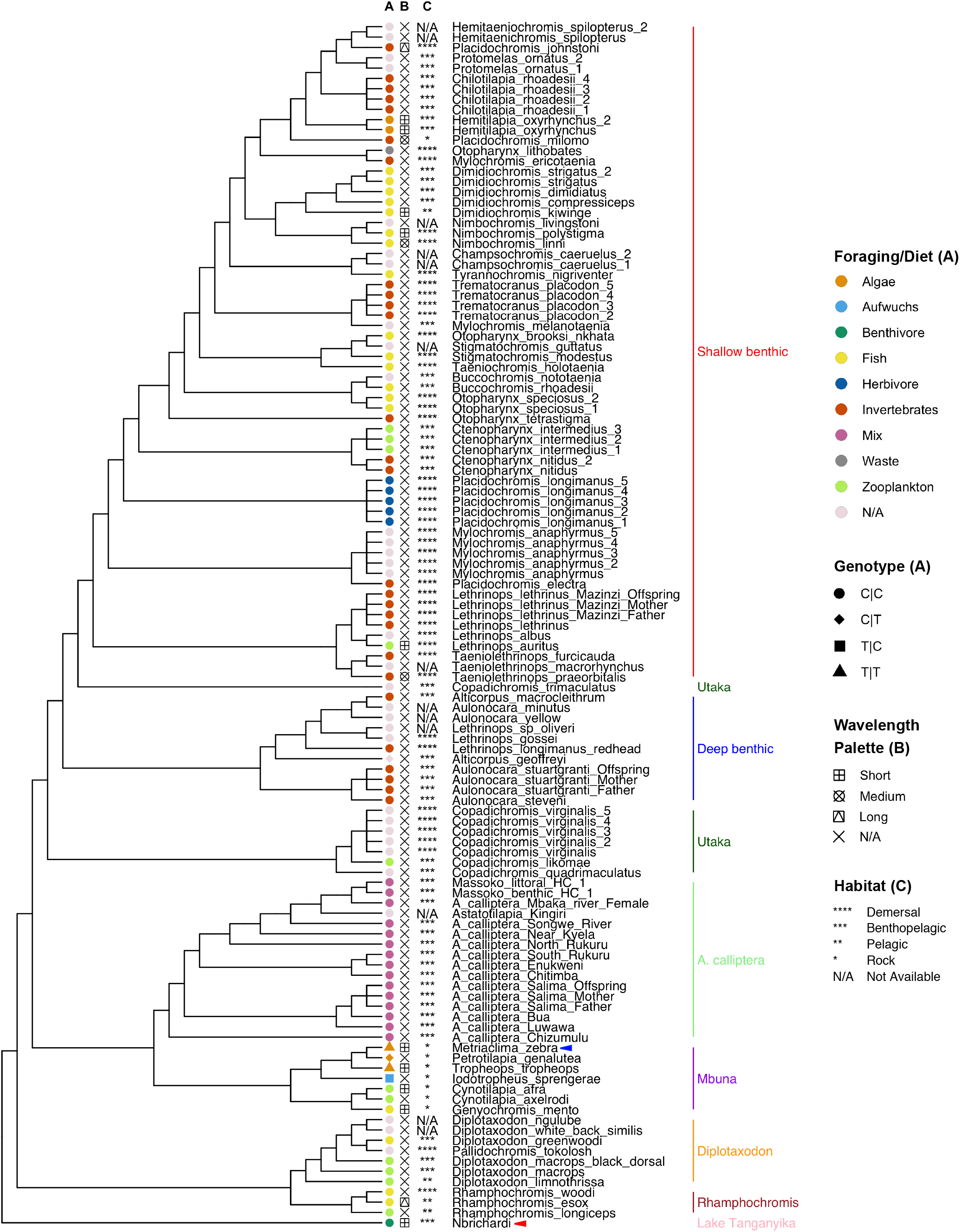
SNP genotypes overlapping NR2C2 TFBS in *M. zebra sws1* promoter and other Lake Malawi species. Lake Malawi phylogeny reproduced from published least controversial and all included species ASTRAL phylogeny [19], including *N. brichardi* as an outgroup. Phylogenetic branches labelled with species sample name (including *M. zebra* with blue arrow and *N. brichardi* with red arrow) and clade according to legends (*right):* A) Species foraging/diet habit (color) [36] and phased SNP genotype (shape) [19]; B) Adult opsin wavelength palette utilized [36]; and C) species habitat [36,80].

## Discussion

The evolutionary ‘tinkering’ of regulatory systems through gene regulatory network divergence can facilitate the evolution of phenotypic diversity and rapid adaptation [38]. Various mechanisms underlie these events, including horizontal gene transfer and regulatory reorganization in bacteria [39]; gene duplication in fungi [40]; *cis*-regulatory expression divergence in flies [41]; variable gene co-expression in worms [42]; dynamic rewiring of TFs in plant leaf shape [11]; coding and non-coding evolution in stickleback fish [43]; alternative splicing [44] and differential rate of gene expression evolution shaped by various selective pressures [45,46] in mammals. However, since very little is known about the combined effect of some of these mechanisms; in-depth analyses of regulatory network evolution can shed light on key contributing mechanisms associated with phenotypic effect across ecologically-diverse species in a phylogeny.

The near 1,500 species of East African cichlid fish, have rapidly radiated and diversified in an explosive manner. Alongside ecological opportunity [17], East African cichlid diversification has been shaped by complex evolutionary and genomic forces, including divergent selection acting upon regulatory regions [18] that is largely based on a canvas of low genetic diversity between species [19]. All of these findings imply the rapid evolution of regulatory networks underlying traits under selection however, little is known about the genome-wide evolution of regulatory networks that may underlie several traits of cichlid phenotypic diversity [47]. Here we developed a novel computational framework to characterize the evolution of regulatory networks and analyse the plausibility of their potential contribution towards phenotypic diversity in closely-related cichlids.

Along the phylogeny, our analyses identified gene co-expression modules with tissuespecific patterns and differential trajectories across six tissues of five cichlids.

Given that the volumes and hence, representation of region-specific cell types of selected organ regions e.g. brain can be different, even between closely related cichlids [48], it is plausible that the observed expression differences between species are driven by changes in cell type abundances. However, given that expression data was generated from the organs of multiple similarly sized adult individuals, and the identification of conserved tissue-specific patterns across all tissues and species e.g. module 1 is eye-specific (Fig .1a), we suspect that the majority of observed coexpression differences are connected to gene regulatory differences. Indeed, these genes are predicted to be regulated by divergent suites of regulators, including TFs that are state-changed in co-expression module assignment. This suggests that in the five cichlids, transcriptional rewiring events and differential gene expression could contribute to phenotypic diversity of the six studied tissues.

*Cis*-regulatory elements (including promoters and enhancers) are central to cichlid gene expression regulation [18], and in this study we show that discrete nucleotide variation at binding sites drives regulatory edge divergence through GRN rewiring events. Our comparative analysis of GRNs across species identifies striking cases of rapid network rewiring for genes known to be involved in traits under natural and/or sexual selection, such as the visual system, possibly shaping cichlid adaptation to a variety of ecological niches. In regulatory regions of visual opsin genes e.g. *sws1*, *in vitro* assays confirm that variations in TFBSs (NR2C2) have driven network structure rewiring between species sharing the same visual palette. Since the modulation of cichlid visual sensitivity occurs through heterochronic shifts in opsin expression [49], our results are consistent with recent findings that visual tuning differences between cichlid species requires regulatory mutations that are constrained by mutational dynamics [50]. Gene duplications have been implicated in cichlid evolutionary divergence, including differences in duplicate TF gene expression [18]. However, due to incomplete lineage sorting and variability in duplicates identified by three separate methods (gene trees, read-depth analyses and array comparative genomic hybridization) [18], we only focus on an example of gene duplication associated with network rewiring. We predict that *rho* is regulated by GATA2, and potentially common to dim-light vision in *M. zebra, A. burtoni* and *O. niloticus* but a duplicate TF, GATA2A, is predicted to be a unique regulator of *rho* in *A. burtoni* and *O. niloticus* only, owing to a variant in the GATA2A TFBS in *M. zebra rho* gene promoter. Furthermore, certain *M. zebra* variants overlapping gene promoter TFBSs e.g. *sws1* (NR2C2) and *rho* (GATA2A) segregate according to phylogeny and ecology of Lake Malawi species [19], suggesting ecotype-associated network rewiring events could be linked to traits under selection in East African cichlid radiations. This is consistent with the adaptive potential of visual system evolution in cichlid species, where changes in spectral tuning of visual signals are likely to lead to dramatic species evolution and possibly speciation events [51]. Given that single regulatory mutations of *Tbx2a* can cause heterochronic shifts in opsin expression and visual tuning diversity between two distinct cichlid species [50], it is likely that the regulatory variation at opsin TFBSs we have predicted and experimentally validated, is a contributing mechanism of evolutionary innovation across many cichlid species. Furthermore, the identification (in predicted TFBSs) of segregating sites across several Lake Malawi species, with conservation of flanking regions, is indicative of shared ancestral variation and functional evolutionary constraint. Beyond the visual systems, we also identify network rewiring of genes associated with several cichlid adaptive traits like, for example, *runx2* associated with jaw morphology [52]; *ednrb1* in pigmentation and egg spots [18,53]; and *egr1* implicated in behavioural phenotypes [54].

The regulatory networks generated here represent a rich scientific resource for the community; powering further molecular analysis of adaptive evolutionary traits in cichlids. As an example, further examination of the vast regulatory factors that we have predicted for the visual systems, that could both up- and down-regulate opsin expression diversity, could further shed light on preliminary studies of *SWS1* [55]*, LWS and RH2* [50] in other cichlid species. This could also involve further functional validation to define a definitive link to trait variation by 1) high-throughput protein-DNA assays to confirm binding of hundreds of sites; 2) reporter and/or cell-based TF-perturbation assays to show that the regulatory variants indeed affect transcription; and 3) genome editing e.g. CRISPR mutations of TFBS variants followed by phenotyping to observe trait effect. Nonetheless, this study is the first genome-wide exploration of GRN evolution in cichlids, and the computational framework (Fig. S-M1) is largely applicable to other phylogenies to study the evolution of GRNs. In this study, we largely focus on *cis*-regulatory mechanisms of GRN rewiring. However, given the potential impact of other genetic mechanisms (protein coding changes, small RNAs, and posttranslational modifications) towards cichlid phenotypic diversity [18,47], our framework can be easily extended to allow for studies on the regulatory effect of other mechanisms e.g. miRNAs, enhancers and extensive gene duplications on network topology during cichlid evolution. Whilst many of the predicted TF-TG relationships could be false positives, our integrative approach ensured that we could apply rigorous filtering at each step; including stringent statistical significance measures, co-expression-based pruning, and all whilst accounting for gene loss and mis-annotations in selected species (see *Methods*). Furthermore, whilst it appears that cichlids utilize an array of regulatory mechanisms that are also shown to drive phenotypic diversity in other organisms [11,40–43,56], we also provide experimental support of selected TF-TG rewiring events in regulatory regions of genes associated with adaptive traits in cichlids [18]. This link between GRN evolution and genes associated with adaptive trait variation in cichlids is further supported by large-scale genotyping studies of the predicted sites in radiating cichlid species [19]; whole genome analysis and transgenesis assays [18]; behavioural and transcriptomic assays [57]; population studies and CRISPR mutant assays [58]; and transcriptomic/*cis*-regulatory assays [34,36,50,55].

## Conclusions

We present a novel approach to reconstruct the evolution of GRNs in vertebrate species. We applied the methods to six tissues from five species that are from the most striking example of a rapid adaptive radiation, the East African cichlids. We build on previous findings, which demonstrate that two contributing factors to rapid cichlid phenotypic diversity are the acceleration of regulatory regions and gene expression divergence. Our novel network-based approach integrates gene co-expression and predicted candidate regulatory regions/variants implicated in the rapid evolution of cichlid traits. We show that tissue-specific gene expression has diverged between five cichlid species and as a case study we presented data from a well-studied trait – the visual system – for which we identified regulatory variation at TFBSs and investigated the plausibility of these variants having an association with the evolution of the visual system. We further demonstrated how the functional disruption of TFBSs indeed abrogates binding of the key regulators, and can drive GRN evolution. Our approach revealed hundreds of novel potential regulatory regions/variants of five cichlid genomes, and our network framework identified several regulatory regions/variants with previously reported associations with a number of known evolutionary traits, thus providing proof-of-concept, as well as novel candidates with no prior known associations. In conclusion, we show that regulatory network evolution can happen rapidly in cichlids, and discrete changes at regulatory binding sites, driving regulatory divergence and network rewiring events are likely to be a contributing source to evolutionary innovations in radiating cichlid species. This approach, with further functional validations, has the potential to identify novel regions linked to other evolutionary traits in cichlids and other evolutionary systems.

## Supporting information

Extended_Data_Table_S-R3A_TFTG1to1_DyNet_rewiring_allfive_species

Extended_Data_Table_S-R3B_Candidate_Genes

Extended_Data_Table_S-R3C_TFTG1to1_DyNet_rewiring_allfive_species_candidates

Extended_Data_Table_S-R3D_TFTGalledges_DyNet_rewiring_allfive_species

Extended_Data_Table_S-R3E_TFTGalledges_DyNet_rewiring_allfive_species_candidates

Extended_Data_Table_S-R3F_TF-TG_evolutionary_rate

## Acknowledgments

We thank the BROAD institute and the Cichlid Genome Consortium for providing full access to genomic data. We thank Dr. Domino Joyce and Prof. Walter Salzburger for providing cichlid species tissues for EMSA validations. TKM, WN, PS, LPD, WH and FDP were supported and the project strategically funded by the Biotechnological and Biosciences Research Council (BBSRC), Institute Strategic Programme BB/J004669/1 and Core Strategic Programme Grants BB/P016774/1; BBS/E/T/000PR9817; and BB/CSP17270/1 at the Earlham Institute. TK was supported by a fellowship in computational biology at Earlham Institute (Norwich, UK) in partnership with the Quadram Institute (Norwich, UK), and strategically funded by BBSRC, UK (BB/J004529/1 and BB/P016774/1). MO was supported by the BBSRC Norwich Research Park Biosciences Doctoral Training Partnership (grant BB/M011216/1). SR was supported by a National Science Foundation (NSF) career award (DBI: 1350677) and with CK and SAK, the McDonnell foundation at The Wisconsin Institute for Discovery.

## Author contributions

CK, SAK and SR constructed gene trees, ran Arboretum and gene ontology (GO) enrichment; TKM, WN and PS developed and ran transcription factor (TF) motif prediction and enrichment; TKM analyzed co-expression modules, enrichment and breadth of gene expression; WN, TKM and WH calculated and analyzed evolutionary rates; SAK and SR generated co-expression edges; TKM reconstructed networks and carried out GO enrichment and analyses; SB and LPD analyzed network structure; MO, TKM and TK analyzed network rewiring; TKM and WN analyzed variants overlapping TFBSs; TKM carried out EMSA; TKM, WH, SR and FDP wrote the manuscript with input from SAK, WN, PS, SB and TK.

## Methods

### A comparative framework to study the evolution of tissue-specific regulatory networks in cichlids

We developed a comparative framework (Fig. S-M1) to infer gene regulatory networks across five representative East African cichlid species - *O. niloticus (On), N. brichardi (Nb), A. burtoni (Ab), P. nyererei (Pn*) and *M. zebra (Mz*). Our framework comprises: (1) identifying modules of co-expressed genes from multi-tissue/multi-species and single-tissue/multi-species data; (2) integrating several datasets (gene expression and *cis*-regulatory regions) to reconstruct gene regulatory networks (GRNs) to find fine-grained tissue-specific network modules; (3) examining factors driving evolutionary innovation in cichlids i.e. nucleotide divergence within regulatory binding sites and determining their mechanistic roles towards regulatory network and module divergence; and (4) using an integration of the reconstructed networks, co-expression modules and enrichment of curated biological processes to interpret GRN evolution of genes in the context of cichlid adaptive traits.

### Inference of multi- and single-tissue transcriptional modules in five cichlids

We ran Arboretum [9], an algorithm for identifying modules of co-expressed genes on gene expression values of six tissues (brain, eye, heart, kidney, muscle, testis) from five cichlid species - *O. niloticus (On), N. brichardi (Nb), A. burtoni (Ab), P. nyererei (Pn*) and *M. zebra (Mz*) [18]. Gene expression values were based on RNA sequencing of several adult individuals; the sample source, tissue isolation, RNA and library preparation, sequencing, assembly and annotation are described previously [18]. To ensure equality in *n*-fold change of expression, the gene expression values were log transformed as: log(x+1), where x is the raw expression value [18], and “log” is the natural logarithm, and then expression was normalized across each gene to have mean zero to be used as input for Arboretum [9]. The *log* expression ratio shown across modules is each genes expression relative to the mean expression across all tissues. Selection of the six tissues allowed us to study tissue-specific associated traits under natural and/or sexual selection in cichlids: Brain (development, behaviour and social interaction); Eye (adaptive water depth/turbidity vision); Heart (blood circulation and stress response); Kidney (haematopoiesis and osmoregulation associated with water adaptation); Muscle (size, shape and movement associated with dimorphism and agility); and Testis (sexual systems associated with behaviour and dimorphism).

In total, 18,799 orthogroups (see Methods: Construction of cichlid gene trees) and their associated expression data and gene tree information were inputted into Arboretum [9]. In total, this represents 59-68% of all protein-coding genes in the five cichlid genomes [18]. Certain annotated cichlid genes could not be included for a few reasons: 1) Lack of tissue expression data for all five species; 2) No mapped reads for selected tissues; 3) Lack of co-expression with other genes; and 4) Use of single development stage (adult). We selected the number of modules using a combination of strategies. First, we tried to identify the optimal number of multi-tissue modules (*k*) automatically from the data by scoring the Arboretum learned model based on the penalized log likelihood. This gave us the optimal *k* of 19, however, we also looked at lower values of k, for example, *k* = 7 and *k* = 15 for examining co-expression clustering patterns in the next step. Second, we manually inspected the modules to see if increases of *k* yield patterns of expression that we have not seen before or generate recurring patterns. Finally, we devised a metric for the top three random initializations, based on a silhouette index, orthology overlap, and cross-species cluster mean dissimilarity; selecting the optimal *k* stable to the initialization. Based on our strategy we found *k =* 10 modules to be optimal. Using a similar approach, this time for single tissues clustering, we found *k =* 5 modules to be optimal.

#### Handling ILS in Arboretum

The Arboretum algorithm internally tries to reconcile a tree that is not obeying the species tree by adding additional duplication and loss events. An alternate approach is to use a different species trees each representing the different ILS types and estimating the parameters of each such tree. However, there are many different cases of ILS, as identified previously [18] and the number of gene trees in each category varied significantly. Estimating the conditional distributions for each branch in each ILS type would not be feasible as there are not enough example trees.

### Cichlid gene trees

By considering the gene tree of 18,799 orthologous groups (orthogroups), Arboretum [9] is able to generate module assignments reflecting many-to-many relationships between orthologs resulting from gene duplication and loss. To construct gene trees with different levels of duplication, we obtained the protein sequences of the longest transcripts from five cichlids as well as stickleback, spotted gar and zebrafish as outgroups. Spotted gar was added as it predates the teleost-specific genome duplication event (3R) and zebrafish, as a model teleost to leverage known molecular interactions as an initial prediction of functional interactions/associations in cichlids based on orthology. We applied OrthoMCL-1.4.0 [59] followed by TreeFix-1.1.10 [60] to learn the reconciled gene trees. We noticed that several of the trees exhibited incomplete lineage sorting (ILS) for the cichlid specific subtree but disappeared once the tree was relearned using the cichlid only species. We therefore relearned gene trees for the cichlid only species - in total, we reconstructed 17858 gene families of which, 108 had gene duplication events. A fraction of these (29 gene families) also exhibited ILS. We also observed ILS for gene groups without gene duplications: of the 17756 gene families that had no duplication, 810 exhibited ILS.

### Functional and transcription factor binding site (TFBS) enrichment in modules

We use the False Discovery Rate (FDR) corrected hypergeometric *P*-value (*q*-value) test to assess enrichment of Gene Ontology (GO) terms and TFBSs (motifs) in a given gene set. In all cases, enrichment is tested using a set-based approach where a set of candidate genes is compared to a background (control set) of either all genes in species modules (18,799 orthogroups) or each genome (stated within figure legend for each test). We summarize the enrichment of terms/motifs with *q*<0.05 statistical significance and conservation in all extant and ancestral species. GO terms for the five cichlids were from those published previously [18]. To study *cis*-regulatory elements likely driving tissue-specific expression patterns, we defined gene promoter regions using up to 5 kb upstream of the transcription start site (TSS) of the gene. This is based on analysing the distribution of motifs in 100nt window regions up to 20kb upstream of each genes TSS, and observing a plateau of motifs (and distribution of CNEs) after ~5kb in each species (Fig. S-M2). Motif enrichment in *cis*-regulatory regions was carried out using TFBSs obtained by the method below, with a background (control set) of all motifs (FDR<0.05) predicted within module gene promoters.

### Transcription factor (TF) motif scanning

TFBSs of known vertebrate transcription factors (TFs) were obtained from the JASPAR vertebrate core motif (2018 release) [61]. Binding peak information from ChIP-seq experiments of various human and mouse TFs were retrieved from GTRD v17.04 [14] and associated to protein coding genes within a vicinity of 10kb. Using core motif sequences available from JASPAR [61] or alternative databases like UniPROBE [62] and HOCOMOCO [63], sequences matching these motifs were identified within the TF binding peaks. In cases where the core motifs were not available for specific TFs with ChIP-seq data, they were predicted *de novo* from the sequences under peaks themselves using MEME [64] with default settings. The aforementioned steps provided a list of transcription factor-target gene (TF-TG) interactions with the exact coordinates of the corresponding binding site(s). Cichlid sites were extrapolated based on 1) gene-level orthology; (based on gene trees above) 2) minimum 70% sequence similarity [65,66] between the vertebrate motif sequence and a sequence within the cichlid promoter; and 3) functional domain overlap as derived using *Interpro scan 5* [67] to both source organisms (human, mouse). Extrapolated sites from the promoters of each cichlid species were used to construct cichlid-species specific (CS) Position Specific Scoring Matrices (PSSMs) for each TF using the *info-gibbs* script from the RSAT tool suite [68]. In cases where the number of extrapolated sites per species was less than three, we aggregated the sites to construct generic cichlid-wide (CW) PSSMs. Using the PSSMs for each TF, we scanned up to 20kb upstream of a genes TSS and conserved noncoding elements (CNEs) with FIMO [69] using either 1) an optimal calculated p-value for each TF PSSM, calculated using the *matrix quality* script from the RSAT tool suite [68], with 1000 matrix permutations; or 2) FIMO [69] default *p-value* (1e-4) for JASPAR [61] PSSMs and PSSMs for which an optimal *p-value* could not be determined. Based on the distribution of motifs in 100nt windows of up to 20kb upstream of gene TSSs (Fig. S-M2), we only retained motifs up to 5kb upstream of a genes TSS as the gene promoter region (Fig. S-M2). Statistically significant motifs were called using a *q-value* (FDR) <0.05 and grouped in confidence levels and scores of: 1a) overlap of mouse and human to cichlid extrapolated - 0.3; 1b) mouse to cichlid extrapolated - 0.2; 1c) human to cichlid extrapolated - 0.15; 2a) FIMO [69] scans using extrapolated CS matrices - 0.125; 2b) FIMO [69] scans using extrapolated CW matrices - 0.110; and 2c) FIMO [69] scans using JASPAR [61] matrices - 0.115. To assess whether motifs are predicted by chance, we also scanned randomised promoter sequences using the same PSSMs.

### Calculating tissue-specificity index (tau)

As a measure for tissue specificity of gene expression, we calculated t (Tau) [70] using log-transformed and normalized gene expression data (as inputted to run Arboretum):

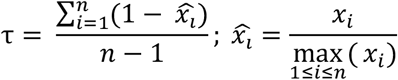

Here, *n* is the number of tissues and *x_i_* is the expression profile component normalized by the maximal component value [70]. The values of tau vary from 0 to 1; ubiquitous or broad expr (t ≤ 0.5); intermediate expr (0.5 < t < 0.9); and tissue-specific or narrow expr (t ≥ 0.9) [70]. Amongst existing methods, t has been shown to be the best for calculating tissue-specificity [71]. Testis normally express far more genes than any other tissue, generally displaying a tissue-specific pattern of expression. As tau was used to assess genome-wide expression levels across all tissues, but between species, testis expression data was included for each species to obtain a true representation of variation in transcriptional programs.

### Variation and evolutionary rate at coding and non-coding regions

We noticed several anomalous start site annotations of genes in *M. zebra, P. nyererei, A. burtoni* and *N. brichardi* when compared to *O. niloticus*. Owing to these anomalies, we re-defined gene start sites to extract putative promoter regions. For each gene, we used the 1^st^ exon (+/- 100bp) of the longest protein-coding sequence in *O. niloticus* to identify, via BLAT-35 [72], corresponding orthologous start sites in the other four cichlid genomes. We filtered the output based on coherent overlap with original annotations [18] and orthogroups in cichlid gene trees. We re-annotated gene start sites (*M. zebra* – 10654/21673; *P. nyererei* – 10030/20611; *A. burtoni* – 10050/23436; *N. brichardi* – 8464/20119) based on BLAT orthology and end sites based on original annotations [18], which was otherwise used for annotating the remaining genes. Based on new annotations, for all 1:1 orthologs where gene expression data is available and there is no overlap of gene bodies, we extracted putative promoter regions, taken as up to 5kb upstream of the transcription start site (TSS) as per methods above. Using *mafft-7.271* [73], we aligned 1:1 orthologous promoter, cds and protein sequences based on orthogrouping in gene trees (see Methods: Construction of cichlid gene trees). We estimated the number of nonsynonymous substitutions per nonsynonymous site (d*N*) and synonymous substitutions per synonymous site (d*S*) in the 1:1 protein alignments using the codeml program in the PAML-4.9 package [74] for each branch and ancestral node in the species tree. Otherwise, we estimated evolutionary rate for each branch and ancestral node in the species tree at promoter regions and fourfold degenerate sites, using 1:1 promoter and cds alignments in baseml and codeml programs in the PAML-4.9 package [74], requiring that at least 10% of the alignment contains nucleotides and that at least 100 nucleotides are present for each species.

By using the published ‘*cichlid-5way.maf*’ [18], we categorised pairwise substitutions for all species and intersected with annotated genomics regions (see Table S-R2a) using *bedtools-2.25.0* intersect [75].

### Reconstructing regulatory networks

To infer essential drivers of tissue-specific expression in cichlids, we constructed regulatory and functional interaction/association networks through the integration of several datasets and approaches (Fig. S-M1). This approach was largely centred on the integration of expression-based and *in-silico* TFBS motif prediction-based networks.

We first used species- and module-specific gene expression levels to infer an expression-based network. For this, we merged the cichlid gene expression data into a single 30 (five species, six tissues) dimensional dataset to learn cichlid-specific transcription factor (TF) - target gene (TG) relationships using the Per Gene Greedy (PGG) approach, a prior expression-based network inference method [24]. We projected the network into species-specific networks by considering edges that would not be present due to gene loss. We then integrated *in-silico* predicted TF-TG edges (see Methods: Transcription Factor motif scanning) based on TFBS predictions in gene promoter regions. To ensure accurate analysis of GRN rewiring through an integrative approach, all collated edges were then pruned to ensure edges were 1) not absent in at least one species due to gene loss/poor annotation; and 2) based on the presence of genes in co-expression modules.

To maintain a structured and connected network approach, we analysed network topology using two methods; firstly, and to ensure suitable integration of co-expression data with all TF-TG predicted edges, one set of all gene nodes and their edges were constrained by Arboretum module assignments to correlate to their respective patterns of tissue-specific expression and co-expression module analysis. Secondly, since all included genes won’t necessarily exhibit tissue-specific co-expression (and cluster accordingly) due to 1) differences in cell type abundance; 2) cell heterogeneity; and 3) small development stage differences, and as well as despite not being co-expressed, the fact that TFs are trans-acting factors able to regulate any gene, we also analysed all network edges for selected candidate genes without constraining based on module assignment (co-expression). Accordingly, for candidate genes with rewired networks, we also analysed network topology without constraining edges based on same module assignment (co-expression) and instead, analysed the Pearson correlation coefficient (PCC) between cross-species TF motif enrichment and tissue expression.

### Functional landscape of reconstructed regulatory networks

We use the FDR-corrected hypergeometric *P*-value to assess enrichment of GO terms for genes in reconstructed networks. We used GO terms for the published five cichlids [18] and carried out enrichment analysis as previously done for Arboretum module genes (see *Methods* above).

### TF-TG gain and loss rates

Gain and loss rate analyses was similar to that performed previously [10]. This approach uses a continuous-time Markov process parameterized by TF-TG edge gain and loss rates, and uses an expectation-maximization (EM) based algorithm to estimate rates [76,77]. The input network comprised target genes of 783 individual regulator genes mapped across the five cichlids species based on gene orthology. Each species regulator required a minimum of 25 edges as <25 edges greatly hinders statistical analysis in this context. This resulted in a total of 345 regulators with 25 to 23,935 edges, with an average of 2,609. Gain and loss rate was estimated for each regulator using the EM-based algorithm on the edge gain and loss pattern across the five cichlid phylogeny. Rates were inferred using published five cichlid branch lengths [18] that described neutral sequence evolution across the species. Stability analysis of rate estimations were carried out as follows: 1) Gain and loss rate input values were scanned from 0-400 in intervals of 5 for each regulator matrix; and 2) From each scan, rates with the greatest likelihood were chosen as the recommended gain and loss rate (<100), defining a final set of inferred rates for 186/345 regulators.

### Regulatory rewiring analysis of gene sets

The DyNet-2.0 package [25], implemented in Cytoscape-3.7.1 [78] was used for network visualization and calculation of dynamic rewiring scores of Transcription Factor (TF) – Target Gene (TG) interactions as predicted by TFBS scanning and TF-TG co-expression relationships (PGG method [24]). To ensure rewiring of TFs are correctly compared between species, and not based on gene loss/poor annotation, we only included edges for analysis where the TF had a 1-to-1 orthologous relationship in species where the TF-TG interaction or non-directed relationship exists. Also, we filtered out any TGs and their TF interaction/relationships if, based on orthologous gene *tblastx* [79], whether the gene was present in the genome but not annotated. Of the 18,799 orthogroups used for generating modules of co-expressed genes and network interactions, 4,209 orthogroups had many-to-many genes actually present in the genome of at least one of the five species. These 4,209 orthogroups were filtered out, retaining 843,168/1,131,812 predicted TF-TG edges across the five species; in summary, these represent edges that are 1) present in at least one species; 2) not absent in at least one species due to gene loss/poor annotation; and 3) based on the presence of nodes in modules of coexpression genes. The 843,168/1,131,812 predicted TF-TG edges across the five species were then used for network rewiring analysis, using the DyNet-2.0 package [25], implemented in Cytoscape-3.7.1 [78], that outputs a degree-corrected rewiring (*D_n_*) score for all orthogroups. The *D_n_* score for each orthogroup was ordered and the mean calculated; the significance of difference of each orthogroups rewiring score against all orthogroups was compared by calculating differences in the standard deviation and applying the non-parametric *Kolmogorov-Smirnov* test (KS-test).

### Identification of segregating sites in TFBSs

Species pairwise variation was identified based on an *M. zebra* v1.1 assembly centred 8way teleost multiz alignment [18]. Pairwise (single-nucleotide) variants were then overlapped with TFBS positions as determined by TF motif scanning using *bedtools-2.25.0* intersect [75]. Pairwise variants of *M.zebra* were overlapped with single nucleotide polymorphisms (SNPs) in Lake Malawi species [19] using *bedtools-2.25.0* intersect [75]. Both sets of pairwise variants overlapping motifs and lake species SNPs were then filtered based on the presence of the same pairwise variant in orthologous promoter alignments. This ensured concordance of whole-genome alignment derived variants with variation in orthologous promoter alignments and predicted motifs. At each step, reference and alternative allele complementation was accounted for to ensure correct overlap. This analysis was not to distinguish population differentiation due to genetic structure, but to instead map regulatory variants onto a number of radiating cichlid species to link to phylogenetic and ecological traits.

### Expression of protein DNA-binding domains (DBDs)

DNA-binding domains (DBDs) of cichlid proteins (NR2C2 and RXRB) were predicted based on alignment and conservation to annotated human and mouse orthologs. *M. zebra* and *N. brichardi* individuals were sacrificed according to schedule 1 killing using overdose of MS-222 (tricaine) at The University of Hull, UK and University of Basel, Switzerland. Tissues were stored in RNA-later using a 1:5 ratio. RNA was extracted from brain, liver and testis tissues of adult *M. zebra* and *N. brichardi* using the RNeasy Plus Mini Kit (Qiagen), achieving RNA integrity (RIN) in the range of 8-10 (Agilent Bioanalyzer Total RNA Pico Assay). First strand cDNA synthesis of DBD-specific regions was carried out using RevertAid H Minus Reverse Transcriptase (Thermo Scientific) and DBDs amplified (2-step RT-PCR) using Platinum Taq DNA Polymerase (Invitrogen) and the primers listed in Table S-M1a. Resulting cDNA was concentrated using Minelute PCR purification (Qiagen) and 700 ng used for *in vitro* transcription/translation using TnT T7 Quick for PCR DNA (Promega) and the Fluorotect GreenLys tRNA (Promega) labelling system. DBD expression was resolved by SDS-PAGE and detection using the fluorescein filter in the ChemiDoc Touch (Bio-Rad) system.

### Electrophoretic mobility shift assay (EMSA) validation of predicted TF-TG interactions

EMSA was carried out using double-stranded Cy5 fluorophore 5’-modified (IDT) DNA probes, *in vitro* expressed DBDs (see above) and the Gel Shift Assay Core System (Promega). Double-stranded DNA probes were generated by annealing sense and antisense oligonucleotides (see Table S-M1a) in annealing buffer (10 mM Tris pH 7.5, 1 mM EDTA, 50 mM NaCl) for 3 mins at 96°C, 1 min at 90°C, 1 min at 85°C, 3 mins at 72°C, 1 min at 65°C, 1 min at 57°C, 1 min at 50°C, 3 mins at 42°C and 3 mins at 25°C in a PCR thermocycler. Binding reactions were carried out in a final volume of 9 μl composed of Gel Shift Binding 5x Buffer (20% glycerol, 5mM MgCl2, 2.5mM EDTA, 2.5mM DTT, 250mM NaCl, 50mM Tris-HCl (pH 7.5), 0.25mg/ml poly(dI-dC)•poly(dI-dC)); 0.01μM of Cy5-dsDNA probe covering the motif and flanking region (28nt); and either 23ng (RXRB, 10.42kDa) or 27ng (NR2C2, 10.73kDa) of expressed DBD. For EMSA validation with increasing Nr2c2 DBD concentrations, 1× = 27ng. For kit controls, 0.01 μM of human SP1 DNA probe was combined with 10,000ng HeLa nuclear extract. Binding reactions were incubated at room temperature for 20 mins. Protein-DNA complexes were resolved on 1mm NuPAGE 4-12% Bis-Tris polyacrylamide gels (Invitrogen) in 0.5× TBE at 100V for 60 mins. Protein-DNA complexes were detected using the Cy5 filter on the ChemiDoc MP (Bio-Rad) system. Exposure settings were adjusted in Image Lab v6.0.1_build34 (Bio-Rad) with same high (5608), low (1152) and gamma (1.0) values set for all associated images.

