## Supplementary Information for "Evolution of regulatory networks associated with traits under selection in cichlids"

#### Results

##### Gene co-expression is tissue-specific and highlights functional evolutionary trajectories

The functional landscape of modules can be related to tissue-specific co-expression (Fig. S-R1b, Ext. Data S-R1A); for example, module 3 with strong brain induction (Fig. 1a) is significantly enriched for neural processes, reflecting core co-regulated networks of genes associated with signal transduction and synaptic activity (FDR <0.05, Fig. S-R1b, Ext. Data S-R1A). Modules with variable tissue co-expression e.g. module 2 (Fig. 1a) have divergent functional enrichment across species, suggesting that proteolysis and ribosomal activity (FDR <0.05, Fig. S-R1b, Ext. Data S-R1A) in kidney and heart physiological function is potentially different among the five cichlid species.

Orthologous genes of each species can be assigned to non-orthologous modules (Fig S-R1a), indicative of potential co-expression divergence and transcriptional rewiring from the LCA (referred to as ‘state changes’ in module assignment). Using a measure of gene expression tissue-specificity, *tau* [1], we show that genes with no state change in module assignment (green bars) have an even, narrow to mid-intermediate breadth of expression whereas state changed genes (red bars) have a narrow to broad expression breadth (Fig. S-R1d), representative of orthologs clustering in non-orthologous modules (state changes). We identified unique state changes and expression divergence of 655 genes along ancestral nodes (Fig. 1b), including several cellular and developmental TFs (51 TFs - Anc4/3; 20 TFs - Anc3/2; 34 TFs - Anc2/1) such as *foxo1, hoxa11* and *lbx1*. These state changed regulatory TFs are also enriched in module gene promoters according to tissue-specific function like, for example, promoters of module 1 genes (eye-specific expression) are significantly enriched (False Discovery Rate, FDR <0.05) for TF motifs involved in retina- and lens-related development/functions e.g. CRX, PITX3 and OTX1 [2] and module 9 genes (brain-specific expression) are significantly enriched (FDR<0.05) for TF motifs involved in brain development/functions e.g. EGR1 [3] and NEUROD2 (Fig. S-R1c, Ext. Data S-R1B). We observe variability in motif enrichment of TFs across species genes e.g. RAR*α/β/γ* and RXR*α/β/γ* [4] of module 1 gene promoters in all species except *N. brichardi* (Fig. S-R1c, Ext. Data S-R1B).

The variability of such motif enrichment, linked to TF expression changes (state-changing) reflects a shifted domain of tissue expression, implying differences in the regulatory control of target genes along the phylogeny. We test this by computing the Pearson correlation coefficient (PCC) between the cross-species TF motif enrichment and tissue-specific expression across species. Our analysis identified several cases of TFs whose expression change/stability was correlated with motif enrichment change. For several TFs that are functionally associated with tissues [5,6], we note a gradual increase in expression along the phylogeny, positively correlated with an increase in motif enrichment e.g. Brain-Cluster2-NFATC3 (PCC 0.99), Testis-Cluster3-LBX1 (PCC 0.98), Kidney-Cluster3-DLX3 (PCC 1.00), Heart-Cluster1-ISL2 (PCC 0.99) (Fig. S-R1e). In other TFs, positive correlation was due to a focused shift in expression where most species have similar fold enrichment e.g. Brain-Cluster2-CDX1 (PCC 0.98) and similar tissue-specific expression profiles (stable within a subset of species), whereas in the divergent species e.g. *N. brichardi*, the expression profile is negatively shifted along with a different motif enrichment (Fig. S-R1e, Extended Data S-R1C-H). There are several cases of this in highly correlated TFs across tissues, examples of which include Brain-EBF1 (PCC 1.0), Eye-E2F7 (PCC 0.97), Heart-TBR1 (PCC 0.99), Kidney-CDX1 (PCC 1.0), Muscle-TBR1 (PCC 1.0) and Testis-EN2 (PCC 0.98) (Fig. S-R1e, Extended Data S-R1C-H).

Overall, we generally note that TFs with a similar motif fold enrichment across all taxa and similar expression across most (but not all) species (state-change in one species) are most positively correlated e.g. Brain-Cluster2-CDX1 (Fig. S-R1e, Extended Data S-R1C).

Similarly, TFs with comparable motif fold enrichment and expression (no state-change) across all species are amongst the most positively correlated e.g. Brain Module9-ZBTB7B, Module9-TEF, Module9-SOX6, Module9-RBPJ, Module9-NFIL3 (Extended Data S-R1C). On the other hand, TFs with subsets of similar motif enrichment in more than one species have slightly reduced, but positively correlated with an expression change in the same species subsets e.g. Eye-Cluster7-RFX4 (PCC 0.95) (Fig. S-R1e, Extended Data S-R1C). At the other end, TFs with no correlation tend to have variability in motif fold enrichment and/or expression (state-change) across all species e.g. Brain Module9-SOX2, Module9-RFX4, Module9-RARG, Module-IRF8, Module9-HSF1. These patterns are present across all tissues (Extended Data S-R1C-H), and therefore shows that there is a reduction in correlation when there are large shifts in motif enrichment and/or expression in several species (several phylogenetic state-changes), but otherwise positively correlated when there are no shifts (no TF state-changes) or subtle shifts (TF state-change in one or subsets of species).

For selected TFs and tissues, the levels of motif enrichment in gene promoters and TF expression are therefore correlated; similar levels of motif enrichment are largely associated with expression conservation (across all species) and subtle expression changes (in one or subsets of species), and therefore more stable (in expression differences) than TFs with variable motif enrichment along the phylogeny. This highlights differential gene regulatory programmes in the five cichlids, that we later confirm to be subtle differences in TFBSs when studying network rewiring events.

Owing to the variability in motif enrichment of retina/lens related TFs of module 1 gene promoters (Fig. S-R1c, see *Main Text*), we test the Pearson correlation coefficient (PCC) between the cross-species TF motif enrichment and eye expression across species. A change in TF motif enrichment between species is representative of either a gain or loss (decay) of TFBSs in eye-expressed (module 1) genes. For module 1 (eye-expressed) gene promoter motifs, we show that some TFs have a near positive correlation (+1) of TF motif enrichment and eye expression e.g. RORC (PCC 0.92), GLI2 (PCC 0.88), CLOCK (PCC 0.86) and CRX (PCC 0.85) across species (Fig. S-R1f *top row*, Extended Data S-R1D). Some of these TFs for example have important functions in modulating opsin expression e.g. CRX [7] and Hedgehog signalling [8] in retinal axon guidance [9] e.g. GLI2. Across these TFs, there are examples in one species (RORC - PCC 0.92; CRX – PCC 0.85) or multiple species e.g. CLOCK (PCC 0.86), where there is an increased enrichment of motifs in module 1 gene promoters (gain of TFBSs) compared to the other species, that are positively correlated with a concurrent increase in TF eye expression (Fig. S-R1f). Therefore, a gain of retinal TF motifs in eye expressed genes is positively correlated with increased expression in the eye. In CRX, a TF known to modulate opsin expression in zebrafish [7] and exhibiting TFBS turnover in cichlid opsin genes [10], there is a significant increase in eye expression (0 > 4+) upon doubling the level of motif enrichment in *M. zebra, P. nyererei, A. burtoni* and *O. niloticus* as compared to *N. brichardi* in general (Fig. S-R1f). This is due to loss (or decay) of retinal motifs associated with decreased eye expression in *N. brichardi* (compared to the ancestral species, *O. niloticus*). However, a similar level of motif enrichment in the haplochromines (*M. zebra, P. nyererei* and *A. burtoni*), that is slightly higher than motif enrichment in *O. niloticus*, is associated with a concurrent higher level of eye expression than *O. niloticus* (Fig. S-R1f). Along the phylogeny, a similar pattern is observed in all TFs with PCC>0.7 (Fig. S-R1f), and indicates that variable motif enrichment in eye-expressed genes is associated with a concurrent change (increase/decrease) in TF eye expression along the phylogeny.

In summary, these correlative patterns of TF expression changes (state-changing module assignment) and TFBS variation, indicative of motif gain and loss, suggest shifted domains of expression in tissues of species along the phylogeny, implying regulatory control by different suites of regulators. This highlights differential gene regulatory programmes, that could be associated with regulatory network changes underpinning traits under selection in cichlids, such as the visual system [11].

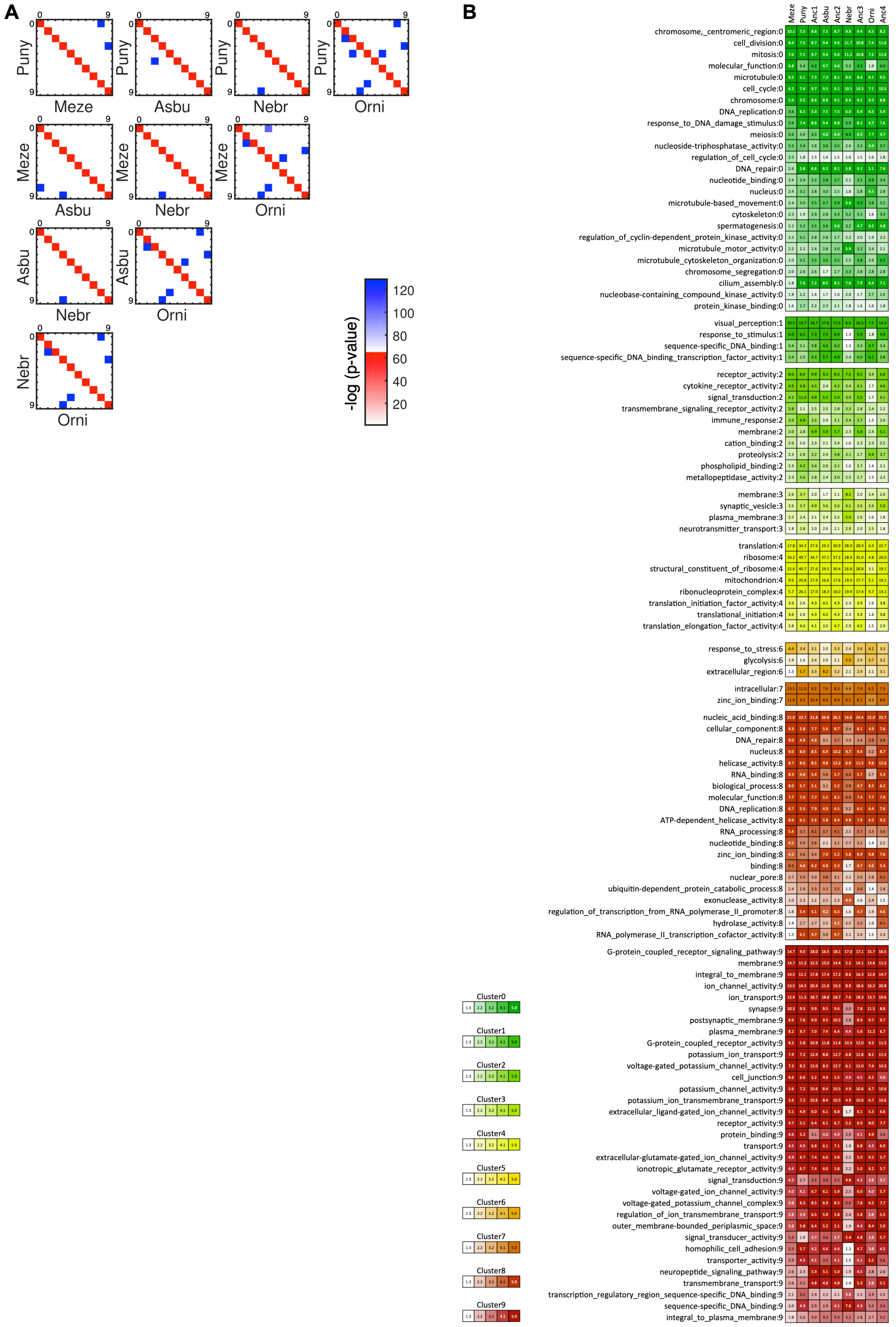


**Supplementary Figure S-R1 – (a) Overlap of module genes between cichlid species.** Shown is the degree overlap of orthologous genes between every module (0-9) pair (rows and columns in each matrix) and in every extant species pair. Diagonal elements (red): overlap between modules of the same ID; off-diagonal elements (blue): overlap between modules of different IDs. Red and blue intensity is proportional to −log (*P*-value) of the hypergeometric distribution (*right,* color scales). **(b) Conserved Gene Ontology (GO) enrichment of modules across all extant and ancestral species.** Conserved enriched terms of significance FDR-corrected *P*-value (*q*-value <0.05) in modules (rows and ':n' module number) are shown for extant and ancestral species (columns) and colored according to module and gradient, -log(*q*-value) in each grid position (see legend, *left*). Set-based hypergeometric test of enrichment carried out using a background of all module genes. Module 5 does not have any conserved enriched terms of significance (FDR<0.05) and instead, all enriched terms for each module are found in Ext. Data S-R1A here: https://figshare.com/s/b0b94f8f108b9a908f53 (http://dx.doi.org/10.6084/m9.figshare.8168246).

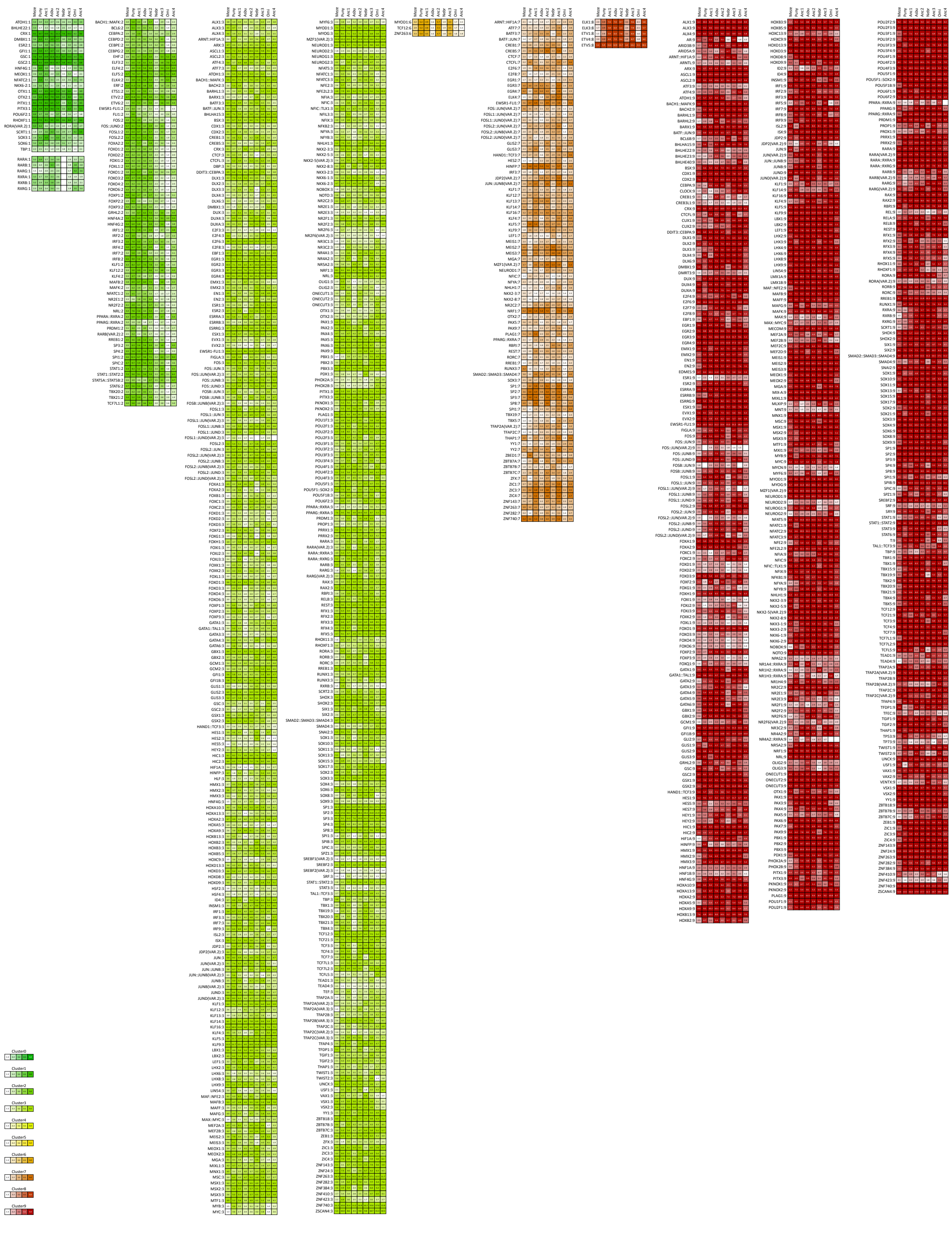

**Supplementary Figure S-R1c – Conserved transcription factor motif enrichment of module gene promoters across all extant and ancestral species.** Conserved enriched motifs of significance FDR-corrected *P*-value (*q*-value <0.05) in modules (rows and ':n' module number) are shown for extant and ancestral species (columns) and colored according to module and gradient, -log(*q*-value) in each grid position (see legend, *left*). All enriched motifs shown are only for conserved across extant and ancestral species modules with the exception of RAR and RXR in module 1 that are shown for the purpose of functional validations. Set-based hypergeometric test of enrichment carried out using a background of all module genes. Selected modules (0, 4 and 5) do not have any conserved enriched motifs of significance (FDR<0.05) and instead, all enriched motifs for each module are found in Ext. Data S-R1B here: https://figshare.com/s/fb59a00eafd3b6c4efa9 (DOI: 10.6084/m9.figshare.8168303).

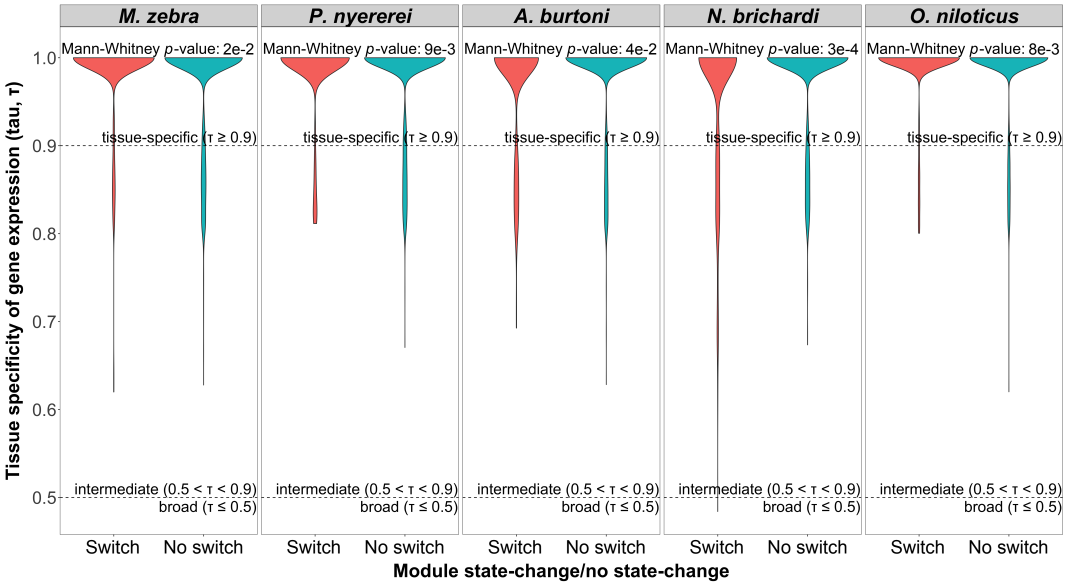


**Supplementary Figure S-R1d – Module genes breadth of expression.** Calculated as Tau (demarcated in plots), shown for each of five species module genes that are switch/state changed (*left*, red violin bars) and no switch/non state changed (*right*, green violin bars). *P* values describing difference between state changed and non-state changed genes breadth of expression calculated using Mann-Whitney test.

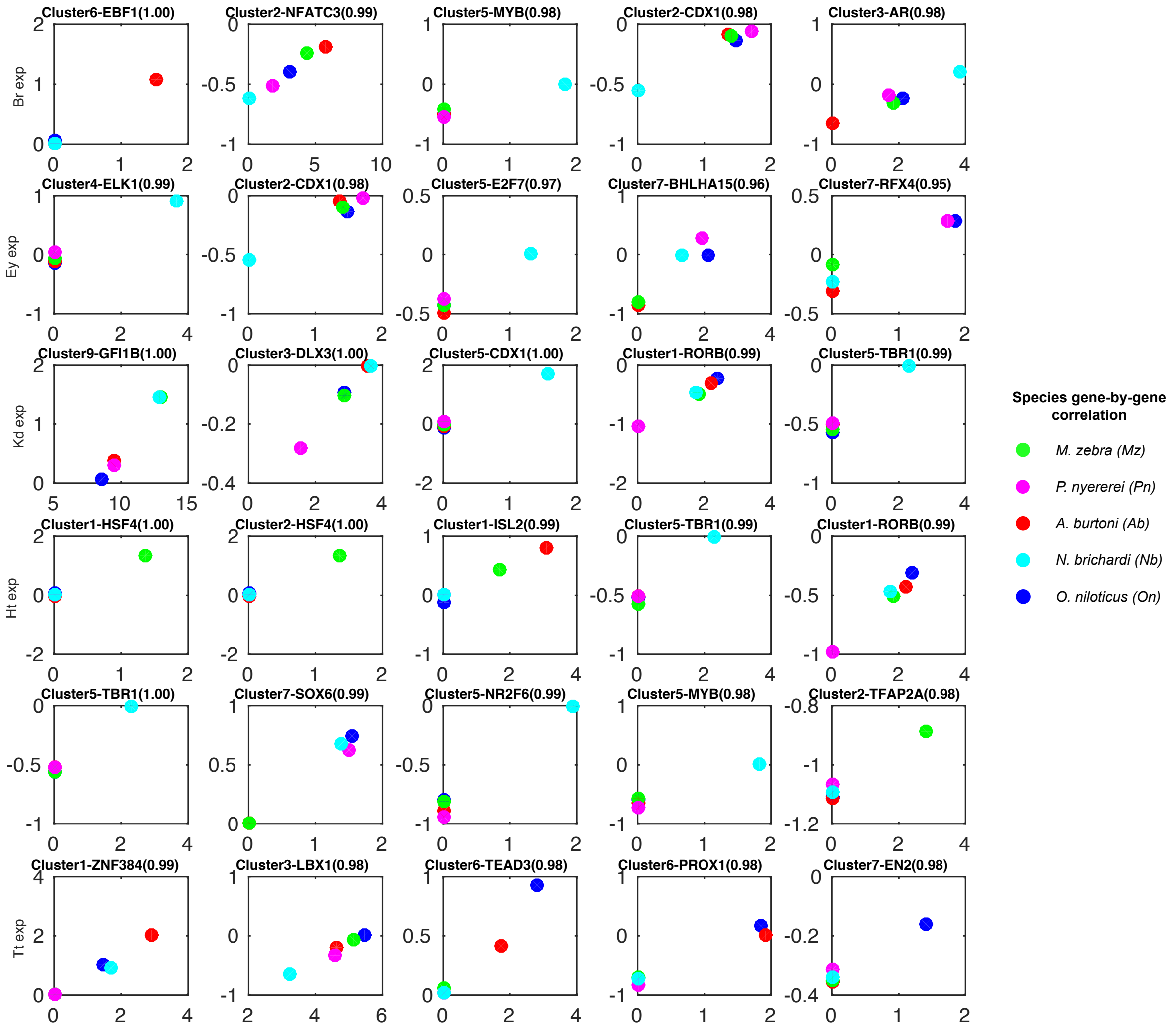


**Supplementary Figure S-R1e – Top five TF motif enrichment and tissue expression correlations across the five cichlid species.** Scatter plots relating to the -log(*p*-value) of fold enrichment (x-axis) and the expression of the transcription factor in six tissues (y-axis). The expression is log zero-mean where the mean of the gene is computed for each species. The title of each scatter plot indicates the module of enrichment (Cluster), TF symbol, and Pearson correlation coefficient (PCC) for all points on the plot. Each dot colour corresponds to a particular species as per legend.

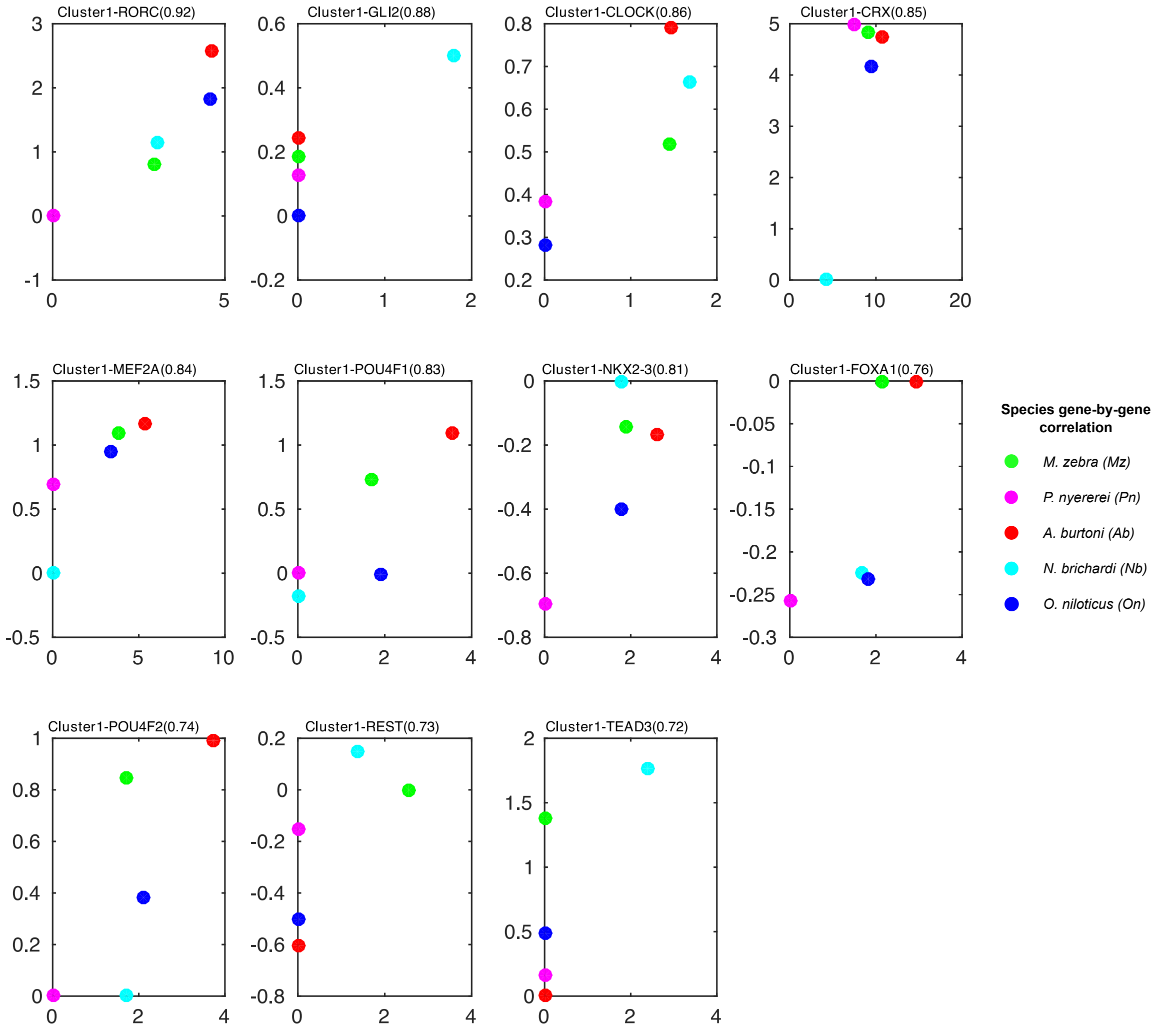


**Supplementary Figure S-R1f – TF motif enrichment in module 1 genes and eye expression correlations >0.7 across the five cichlid species.** Scatter plots relating to the -log(*p*-value) of fold enrichment (x-axis) and the expression of the transcription factor in eye tissue (y-axis). The expression is log zero-mean where the mean of the gene is computed for each species. The title of each scatter plot indicates the module of enrichment (Cluster1), TF symbol, and Pearson correlation coefficient (PCC) for all points on the plot. Each dot colour corresponds to a particular species as per legend.

#### Fine scale nucleotide variation at TF binding sites drives functional regulatory divergence in cichlids through GRN rewiring

The impact of noncoding sequence variation on gene expression was tested based on the evolutionary rate of 4622 1:1 orthologous gene promoter sequences against synonymous (fourfold degenerate) sites of protein coding regions, used as a proxy for neutral evolution. In the five cichlid genomes, there is no significant increase in evolutionary rate at promoter regions compared to fourfold-degenerate sites (Fig. S-R2aA, C, E, and F), with also no difference in promoter evolutionary rate between state-changed and non-state changed genes. We identify very few outlier genes with significantly higher evolutionary rate at promoter regions than corresponding fourfold sites at ancestral nodes (12-351 genes, Fig. S-R2aB) and within species (29-352 genes, Fig. S-R2aD), indicative of small-scale changes in promoter regions. Given the lack of significant evolutionary rate in the majority of gene promoter regions, we hypothesize that discrete changes that could otherwise alter *cis*-regulatory binding sites, could drive gene expression variation in the five cichlids.

Owing to the discrete nucleotide variations observed in various regulatory regions, including selected promoter regions (Fig. S-R2a), we expect that some of the variation may occur at TFBSs (Fig. S-R1c). We identified several pairwise variants between the five cichlids that overlap various genomic regions (Table S-R2a), including state-changed and non-state changed gene promoters and 3' UTRs (Fig. S-R2b). A large proportion of pairwise species variants (12 to 25 million) overlap predicted TFBSs in promoter regions, constituting 14-22% of all pairwise variants in the five species (Table S-R2a, Fig. S-R2b). GO enrichment analysis of cichlid pairwise variants overlapping gene regulatory regions highlight associations with key molecular processes e.g. signal transduction - non-state changed promoter TFBSs (Fig. S-R2c). These findings imply that discrete nucleotide variation at regulatory binding sites could drive functional gene co-expression variation in cichlids through GRN rewiring events.

We focused on regulatory interactions with DNA, most prominently the identification and analysis across species of TF binding to gene promoters (Table S-R3a) as 1) a large proportion of all pairwise species variation (14-22%) overlap TFBSs (Table S-R2a, Fig. S-R2b) and hence, disrupted binding sites will offer insights into GRN rewiring between species; 2) gene orthology is well characterized (as opposed to other regulators, like miRNAs); and 3) direct correlation to tissue co-expression patterns can be made (Fig. 1a). We used a few metrics to study large-scale network rewiring between species, including the analyses of 1) state changes in module assignment; and 2) rewired network edges based on DyNet[12] network rewiring scores (see *Methods*). We first focused on 6,844 1-to-1 orthologous genes in 215,810 TF-TG interactions, termed 'TF-TG 1-to-1 edges', along the five cichlid tree. In total, we identify 4,060-9,423/215,810 TF-TG 1-to-1 edges that are rewired (in a focal vs other species) along the cichlid tree (FDR<0.05, Fig. 2a), and linked to module assignment state changes of 50-70 out of 379 TFs. Given that the level of statistical significance applied (FDR<0.05) could include all 4,060-9,423 (2-4%) rewired and TF state-changed edges, we further analysed the drop-out over more stringent thresholds. Analyses at more stringent thresholds (than FDR<0.05) maintain a similar number of rewired edges ranging from around 1% retained (FDR<0.01) to 2.3% retained (FDR<0.04) and thus, very few rewired edges are likely to be false positives. In the 215,810 TF-TG 1-to-1 edges, we identify 31 out of 90 teleost and cichlid trait genes associated with morphogenesis from previous studies (Extended Data Table S-R3B) that have rewired GRNs based on their DyNet [12] degree-corrected rewiring ($D_n$) score (Extended Data Table S-R3A). A total of 9 out of 31 morphogenesis genes have a few standard deviations higher degree-corrected rewiring ($D_n$) score than the mean (0.17±0.03 SD) score of all 1-to-1 orthologs (Fig. 2c – left violin plot, orange dots; Extended Data Table S-R3C). Furthermore, the degree-corrected rewiring ($D_n$) score of these nine genes (Fig. 2c – left violin plot, orange dots) is significantly higher (Kolmogorov–Smirnov KS-test *p-value* = 0.0006) and thus, exhibit more rewired edges compared to rewired 1-to-1 ortholog edges (Fig. 2c – left violin plot, black dots). Examples of the nine genes include *gdf10b* associated with axonal outgrowth and fast evolving in cichlids [13]; *rh2* – a visual opsin gene[11]; *draxin* – a neural development gene under selection in deepwater cichlid species [14]; and *cntn4,* also associated with neural development and fast evolving in cichlids [13] (Fig. 2c – left violin plot; Extended Data Table S-R3C). To also study rewired networks of orthologs not shared along each taxon of the five cichlid tree, we extended our analyses beyond focusing on 6,844 1-to-1 orthologs only, by also including 7,746 many-to-many orthogroups (see *Methods*) in a set of 843,168 'TF-TG all edges' across the five species. In these edges, we identify 89 out of 90 teleost and cichlid trait genes associated with morphogenesis from previous studies (Extended Data Table S-R3B) that have rewired GRNs based on their DyNet [12] degree-corrected rewiring ($D_n$) score (Extended Data Table S-R3A). A total of 60 out of the 89 morphogenesis genes have a few standard deviations higher degree-corrected rewiring ($D_n$) score than the mean (0.23±0.007 SD) score of all orthologs (Fig. 2c – right violin plot, orange dots; Extended Data Table S-R3E). Furthermore, the degree-corrected rewiring ($D_n$) score of these 60 genes (Fig. 2c – right violin plot, orange dots) is significantly different (KS-test *p-value* = 6e-14) and thus, exhibit more rewired edges compared to the rewiring of all ortholog edges (Fig. 2c – right violin plot, black dots). These genes include most visual opsins e.g. *rho, sws2* and *sws1* [11]; genes associated with photoreceptor cell differentiation, *actr1b*[15]; eye development, *pax6a* [2]; and neuro- and retino- genesis, *irx1* [16,17] (Fig. 2c – right violin plot; Extended Data Table S-R3E).

Since cichlid adaptive radiations extend far beyond the five species we study here, we extended our analyses to include radiating lake species data. In this analysis, we link to previous studies and resources made available [18] to genotype our variants and study how they segregate in the Lake Malawi phylogeny. We overlapped all identified TFBS variants between *M. zebra* (a Lake Malawi species) and the other four cichlids, onto corresponding positions of variants identified in a 73 Lake Malawi species (134 individuals) genome alignment [18]. Using *M. zebra* genotypes as a reference, the expectation would be that more variants would exist with different lake species (like *N. brichardi* from Lake Tanganyika) as well as distantly related same lake species (from Lake Malawi), than closely related same lake species. Of the total 5710 identified variants, the mean number of different genotypes at corresponding positions (vs *M. zebra*) is higher at distant (Lake Tanganyika – 4278; Rhamphochromis – 1756; Diplotaxadon – 1968; Shallow benthic – 1758) than at closely related (*A. calliptera* – 1674) or same (Mbuna – 1540) clade species (Fig. S-R3b). This analysis formed the basis for focusing on particular variants that can be associated with traits under selection e.g. visual systems[11] (*sws1)* and morphogenesis[13] (*cntn4*); to ultimately study variants in TFBSs that segregate according to phylogeny and ecology of radiating lake species.

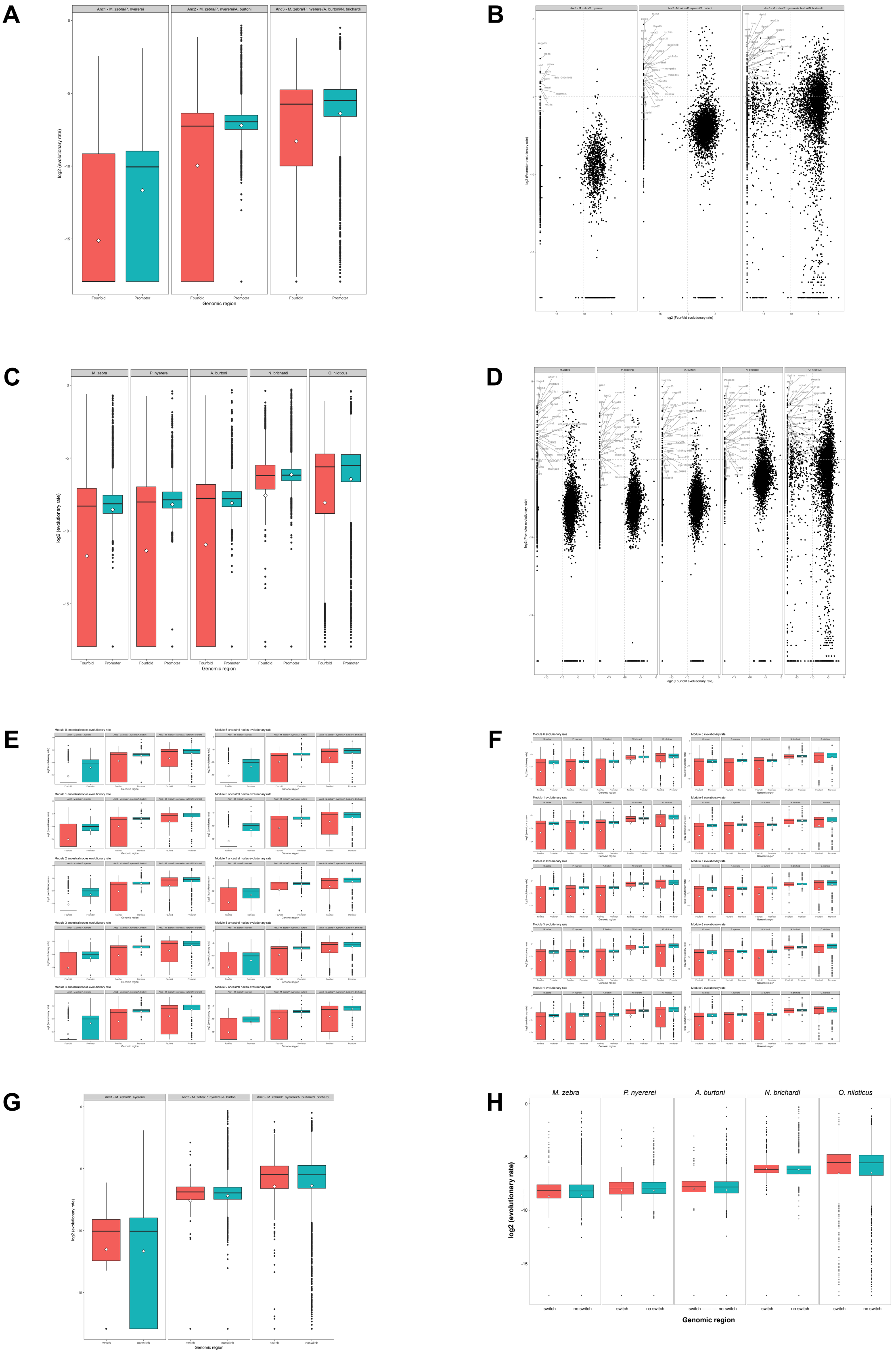


**Supplementary Figure S-R2a – Evolutionary rate at cichlid promoter regions. (A)**
Boxplot of *log2* evolutionary rate in promoters (green bars, *right*) and fourfold degenerate

sites (red bars, *left*) of 1:1 orthologous cichlid genes at each ancestral node; **(B)** Dot plot of evolutionary rates at promoter and fourfold degenerate sites of 1:1 orthologous cichlid genes at each ancestral node. Boundaries (grey dotted line) and top 30 outliers genes with high log2 promoter evolutionary (>-5) and low log2 fourfold site rate (<-10) are marked within; **(C)** Boxplot of *log2* evolutionary rate in promoters (green bars, *right*) and fourfold degenerate sites (red bars, *left*) of 1:1 orthologous cichlid genes at each branch; **(D)** Dot plot of evolutionary rates at promoter and fourfold degenerate sites of 1:1 orthologous cichlid genes at each branch. Boundaries (grey dotted line) and top 30 outliers genes with high log2 promoter evolutionary (>-5) and low log2 fourfold site rate (<-10) are marked within; **(E)** Boxplot of *log2* evolutionary rate in promoters (green bars, *right*) and fourfold degenerate sites (red bars, *left*) of co-expressed 1:1 orthologous cichlid genes at each ancestral node; **(F)** Boxplot of *log2* evolutionary rate in promoters (green bars, *right*) and fourfold degenerate sites (red bars, *left*) of co-expressed 1:1 orthologous cichlid genes at each branch; **(G)** Boxplot of *log2* evolutionary rate in promoters regions of state-changed/switching (red bars, *left*) and non-state changed/non-switched (green bars, *right*) 1:1 orthologous cichlid genes at each ancestral node (switches against LCA, as in Fig. R1B); **(H)** Boxplot of *log2* evolutionary rate in promoters regions of state-changed/switching (red bars, *left*) and non-state changed/non-switched (green bars, *right*) 1:1 orthologous cichlid genes at each branch (state changes against all other species, as in Fig. 1B).

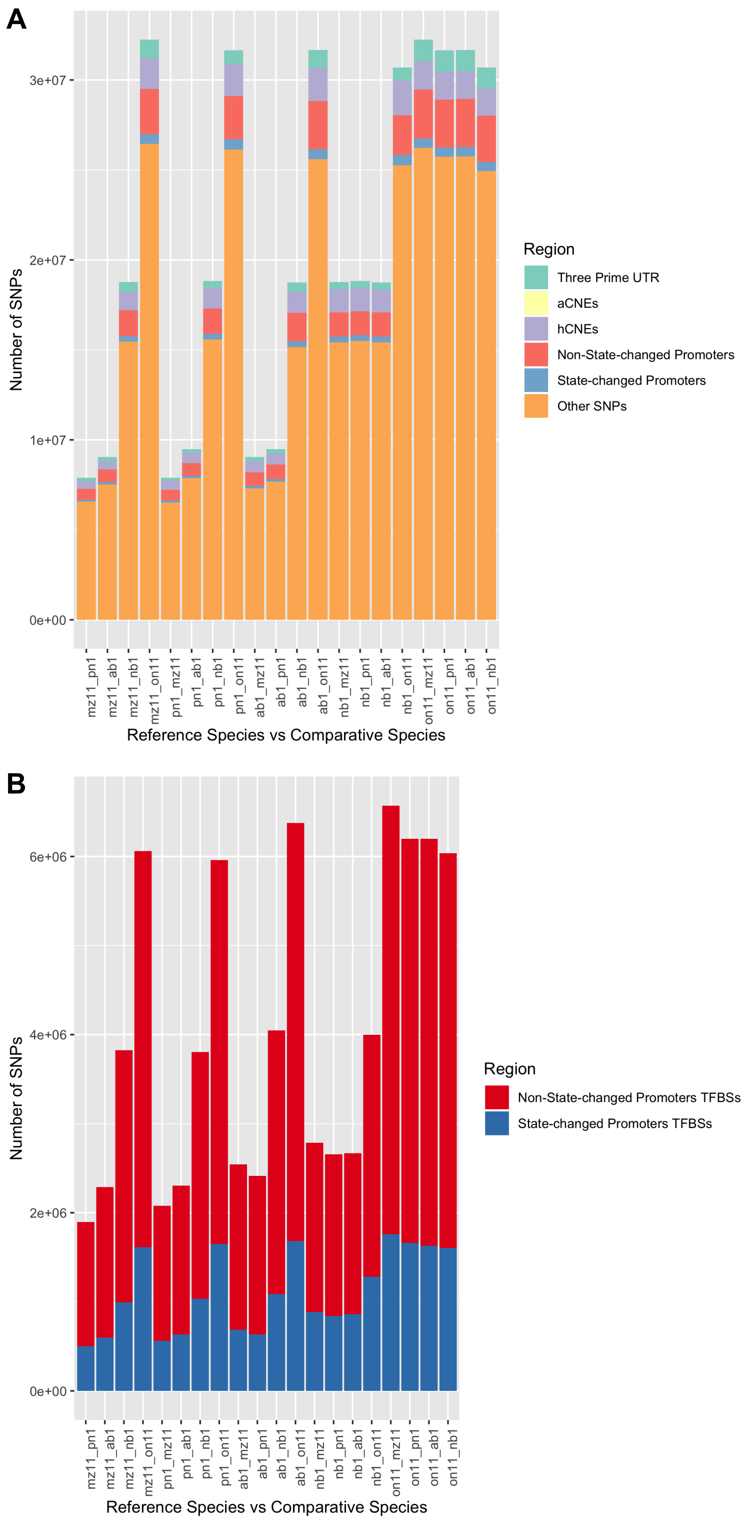


**Supplementary Figure S-R2b – Pairwise variants overlapping various regulatory regions between the five cichlids.** Number of variants (y-axis) overlapping regulatory regions (colored bars) derived from pairwise comparisons (x-axis). Species are named accordingly: *M. zebra* (mz11); *P. nyererei* (pn1); *A. burtoni* (ab1); *N. brichardi* (nb1); and *O. niloticus* (on11).

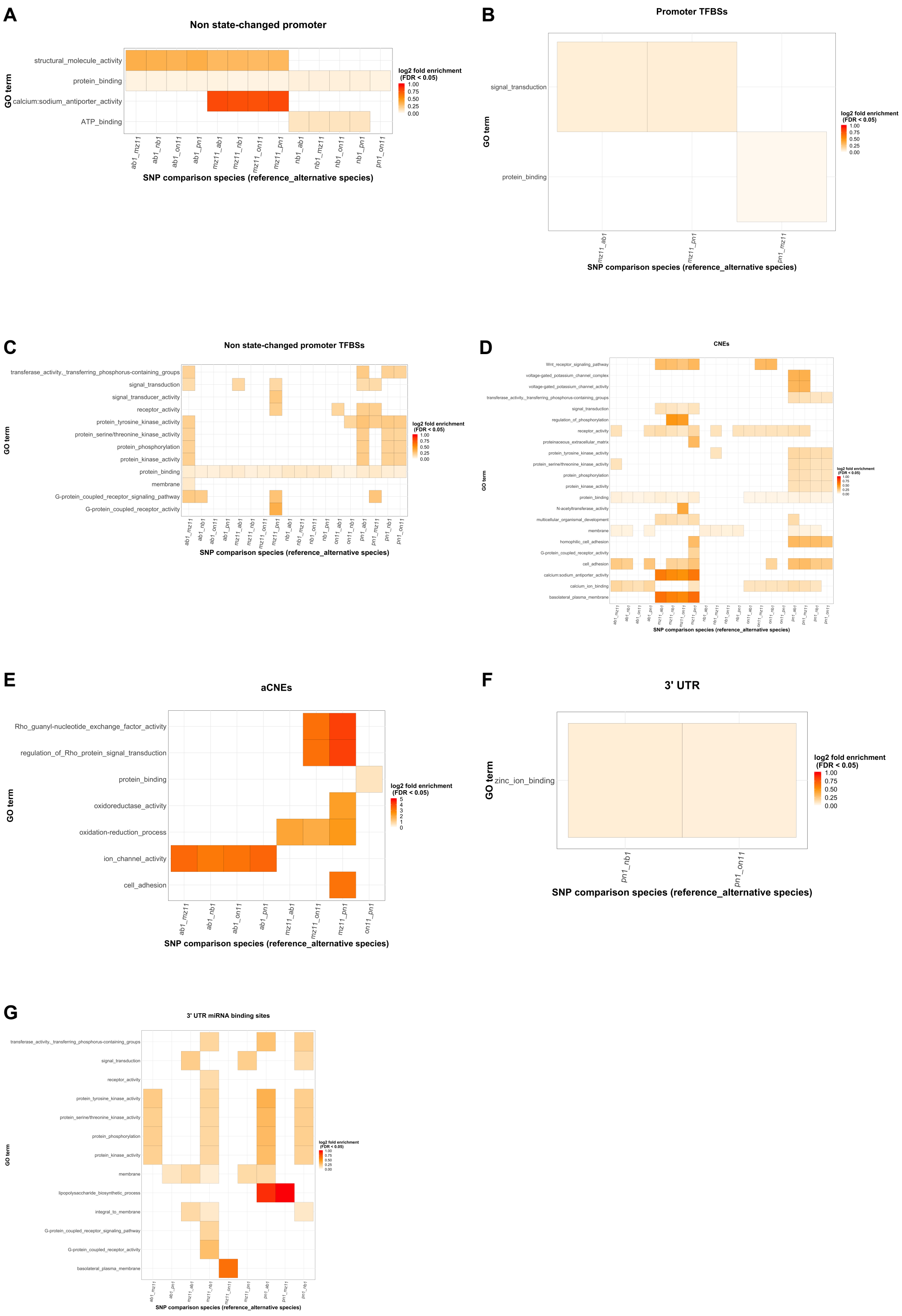


**Supplementary Figure S-R2c – Gene Ontology (GO) enrichment of pairwise variants overlapping various regulatory regions between the five cichlids.** Enriched terms (y-axis) shown as grid heatmap of *log10* fold enrichment (legend on *right*, FDR <0.05) of pairwise species comparisons (x-axis). Set-based hypergeometric test of enrichment carried out using a background of all genes in each species genome. No significant (FDR< 0.05) enrichment in ‘state-changed promoters’ and ‘state-changed promoter TFBSs’.

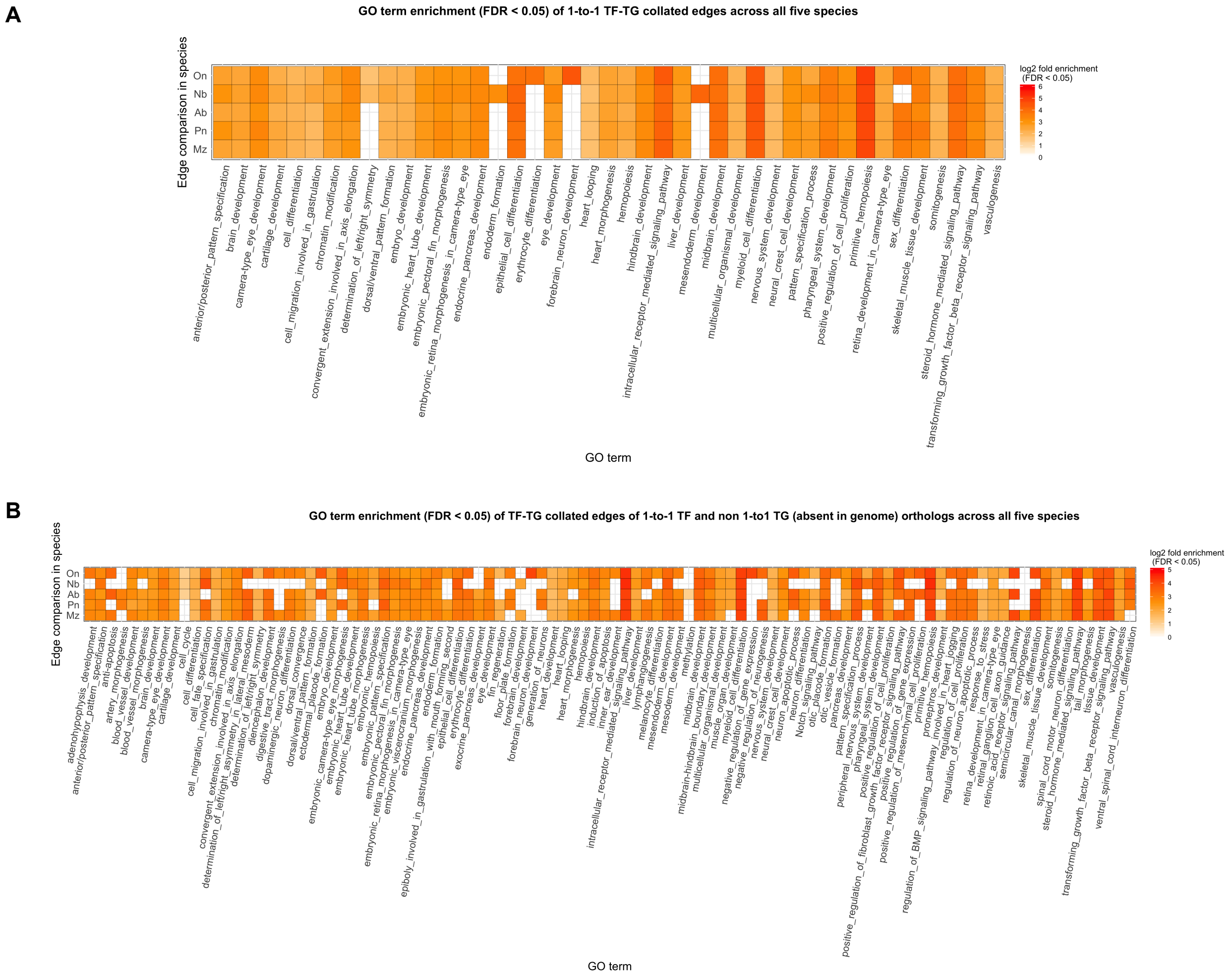


**Supplementary Figure S-R3a - Gene Ontology (GO) enrichment of collated network edges**. GO enrichment shown for both transcription factors (TF) and target genes (TG) in network edges of **(A)** 1-to-1 TF-TG; and **(B)** 1-to-1 TF and non-1-to-1 TG (absent in genome) orthologs in all five species. Species (y-axis) and significant FDR-corrected *P*-value (*q*-value <0.05) GO terms (x-axis) where *log10* fold enrichment shown as grid heatmap (legend on *right*). Set-based hypergeometric test of enrichment carried out using a background set of all module genes (18,799 orthogroups).

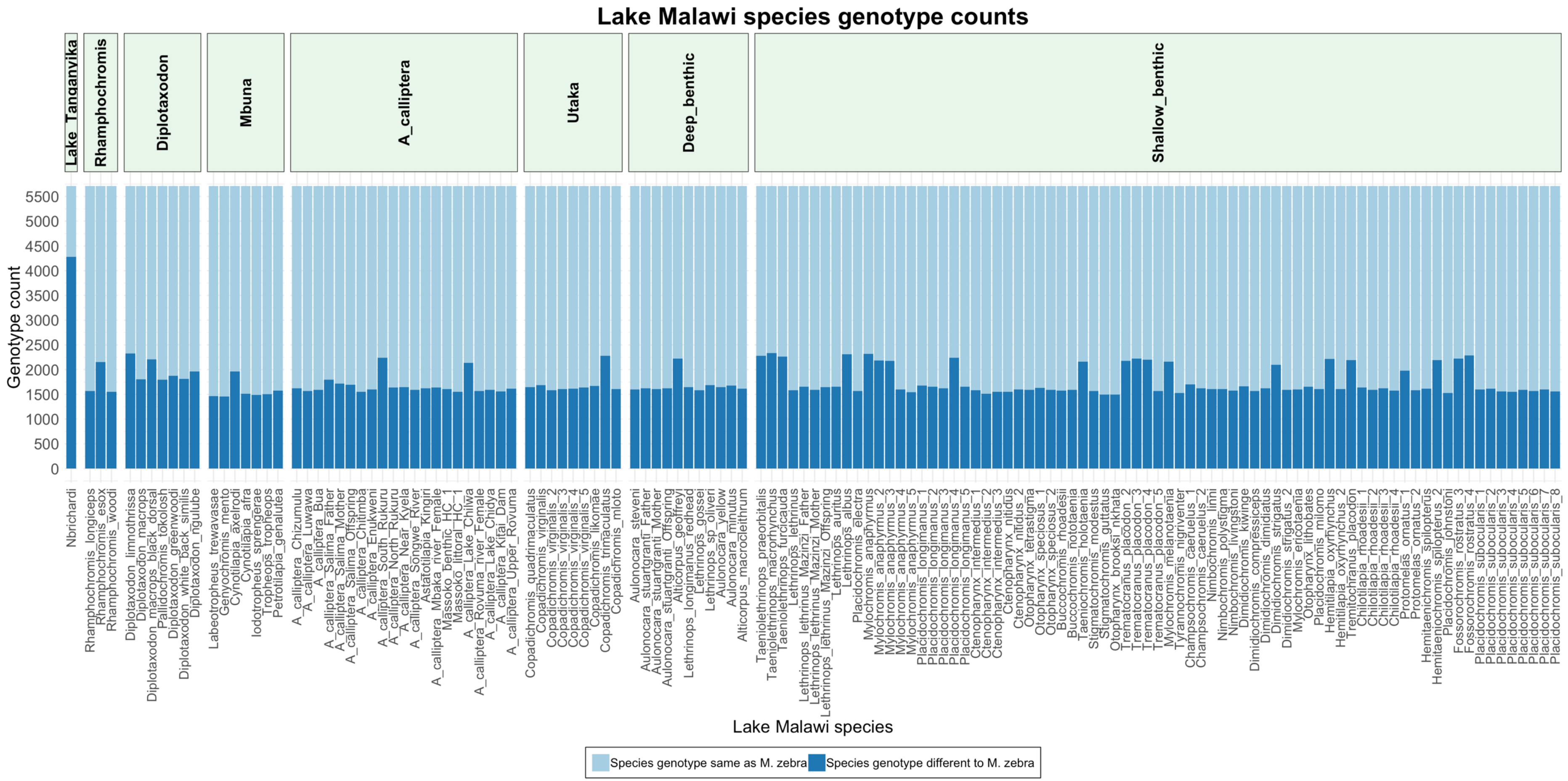


**Supplementary Figure S-R3b - Pairwise variants in candidate gene promoter TFBSs that segregate or are conserved in *M. zebra* and Lake Malawi species.** Genotype counts (y-axis) of conserved (light blue bars) and diverged (dark blue bars) genotypes compared to *M. zebra* in Lake Malawi species (x-axis). Lake Malawi species are ordered and placed into clades according to the published ASTRAL phylogeny; the least controversial and lowest mean distance phylogeny that included all sampled species[18].

#### *Cis*-regulatory changes lead to GRN alterations that control traits linked to phylogeny and ecology of East African cichlid radiations

*Sws1* (ultraviolet) opsin is utilized as part of the short-wavelength sensitive palette in *N. brichardi* and *M. zebra*. A comparison of the *sws1* network of both species identifies several common (18 TFs) and unique (*N. brichardi* – 6 TFs; *M. zebra* – 38 TFs) regulators (Fig. 3a). By ranking and plotting the significance (FDR<0.05) of the unique TF-*sws1* edges, we identify that there is a larger proportion of more significant unique regulators of *sws1* in *M. zebra* (18/38 TFs, orange dots less than mean, Fig. 3a *bottom right*) than *N. brichardi* (2/6 TFs, orange dots less than mean, Fig. 3a *bottom right*).

We focus on NR2C2 and RXRB owing to the significance of this unique predicted *sws1* edge interaction in *N. brichardi* (Fig. 3a, *bottom right*), and show that a candidate polymorphic site in *M. zebra sws1* gene promoter has likely disrupted binding of NR2C2 (Fig. 3). The candidate variant that has likely disrupted binding of NR2C2, and possibly regulation of *M. zebra sws1* (Fig. 3) is an outlier homozygous SNP (A|A) when compared to all other four species (*P. nyererei, A. burtoni, N. brichardi* and *O. niloticus*) that have the homozygous G/G genotype (negative orientation). The outlier homozygous SNP in *M. zebra* as well as flanking sequence predicted as the *M. zebra sws1* gene promoter and used for EMSA validation (Fig. 3c-d) is 100% conserved in the recently published chromosome-scale *M. zebra* assembly [19,20].

We accept that the EMSA binding validation (Fig. 3) does not provide evidence for disrupted regulation of *sws1* alone*.* To address the point of whether NR2C2 are contributing towards regulatory network rewiring, we sought to use our expression data to first predict the regulators for *sws1* by using a regression model. For this, we used expression data from all tissues and species to predict the regulatory relationship based on the co-variation of a TF's

expression level with a TG's expression level across tissues and species. Using this we established a "skeleton" network that predicts the potential regulators of *sws1*; among the top regulators of *sws1* were vision-related regulators such as VSX2[21] and NRL[22].
We next compared these predicted expression-based regulators to see the overlap with the motif based (*cis*) regulators however, there was low overlap between the different networks. This is not surprising and we would expect improved overlap upon the inclusion of more tissues (for expression data) and further species-specific motif information derived from epigenetic techniques e.g. ChIP-seq. The regulators that did overlap however, were among the top regulators, namely, TBX4 (confidence 0.9). We next assessed the correlation of the top regulators predicted by expression, including TBX4, NRL, NR2C2 and RXRB to the expression of *sws1* based on three criteria: (a) global correlation (gcc) using all tissues and species (Fig. S-R4eA *on left*, Fig. S-R4eB – first column, *gcc*), (b) tissue-specific correlation asking to what extent these dependencies are preserved based on eye-specific correlation (Fig. S-R4eA *on right*, Fig. S-R4eB), (c) species-specific correlation (Fig. S-R4eC). In the tissue-specific expression, we can think of each species as a pseudo knockdown/upregulation experiment of the regulator. If the cross-species variation of expression of the regulator is predictive of the variation in the target, we can conclude that the edge might be rewired because the strength of regulation varies. Based on global correlations, we see that TBX4 and NRL are well-correlated to the expression of *sws1*. Both NR2C2 (and RXRB) have relatively lower correlation, but they rank comparably to the expression-based regulators (Fig. S-R4eA *on left*, Fig. S-R4eB – first column, *gcc*), and can therefore explain some of the variation of *sws1*. Focusing on eye-specific correlation only (Fig. S-R4eA *on right*, Fig. S-R4eB – second column, *cc*), we see that the expression of TBX4 across species is predictive *of sws1* expression. NRL is not as predictive, largely due to relatively lower expression in N. brichardi, despite a high expression of *sws1*. Whilst both NR2C2 and RXRB are negatively correlated in the eye, NR2C2's profile is more correlated compared to RXRB – supportive of our functional validations (Fig. 3). Finally, focusing within each species (Fig. S-R4eC), we find that RXRB's expression is not predictive *of sws1* in N.

brichardi (Pearson CC=~0), but is predictive in *M. zebra* (Pearson CC=0.46). In contrast, NR2C2 is predictive *of sws1* expression in both *M. zebra* (Pearson CC=0.30) and *N. brichardi* (Pearson CC=0.27). This suggests that, in *M. zebra*, based on expression either NR2C2 or RXRB could regulate *sws1* but the correlation is weak, whilst RXRB's correlation is slightly higher. However, in *N. brichardi*, only NR2C2 could to regulate *sws1*, supportive of our functional validations (Fig. 3).

We ran a similar analysis for the *rho* target gene, considering CRX and VSX2 as the top expression regulators, and the duplicated TFs, GATA2A and GATA2 (Fig. S-R4f) that are predicted to regulate in selected species (see *Main Text*). Based on global correlations, both GATA2A and GATA2 rank among the expression-based regulators with positive correlation (Fig. S-R4fA) and thus, have the potential to regulate the expression of *rho*. Focusing on the correlations in eye, we see that whilst both GATA2A and GATA2 have a negative correlation, GATA2A ranks better than GATA2 (Fig. S-R4fA, Fig. S-R4fB – second column, *cc*). The species-specific correlations are most informative for this regulatory edge (Fig. S-R4fC). We find that in *O. niloticus* and *A. burtoni,* GATA2A is positively correlated (0.79 and 0.21, respectively), however, in *M. zebra*, where the GATA2A edge is lost (Fig. S-R4c), GATA2A has a negative correlation. GATA2 is still positively correlated (Fig. S-R4c), supportive of its predicted regulation of *rho* in *M. zebra, A. burtoni* and *O. niloticus* (Fig. S-R4c). These species-specific correlations are therefore supportive of GATA2's possible conserved role in all three species, while a more divergent role and binding (Fig. 4) of GATA2A.

To summarise, our expression analysis suggests that the functional binding validations of selected *cis*-regulators, NR2C2 and GATA2A rank comparably among the expression-based regulators and therefore have regulatory potential. NR2C2 is more likely to regulate *sws1 in N. brichardi* compared to RXRB, while both could be plausible in *M. zebra* but still weakly correlated. Given the role of NR2C2 in nuclear receptor signalling [23], important for eye development/function and ability to enhance or repress gene expression in response to environmental cues [24], we suspect an important role in opsin gene expression and cichlid visual system adaptation. GATA2 exhibits a conserved positive correlation with *rho* in all three species with *rho* expression, however, GATA2A's correlation with *rho* in *M. zebra* is significantly lower than in *A. burtoni* and *O. niloticus*, suggesting it is likely not regulating *rho* in *M. zebra*. Taken together these results leverage the natural variation in expression across species and tissues to simulate a perturbation experiment and provide expression-based evidence of the regulatory connections of NR2C2>*sws1,* GATA2A>*rho* and GATA2>*rho*.

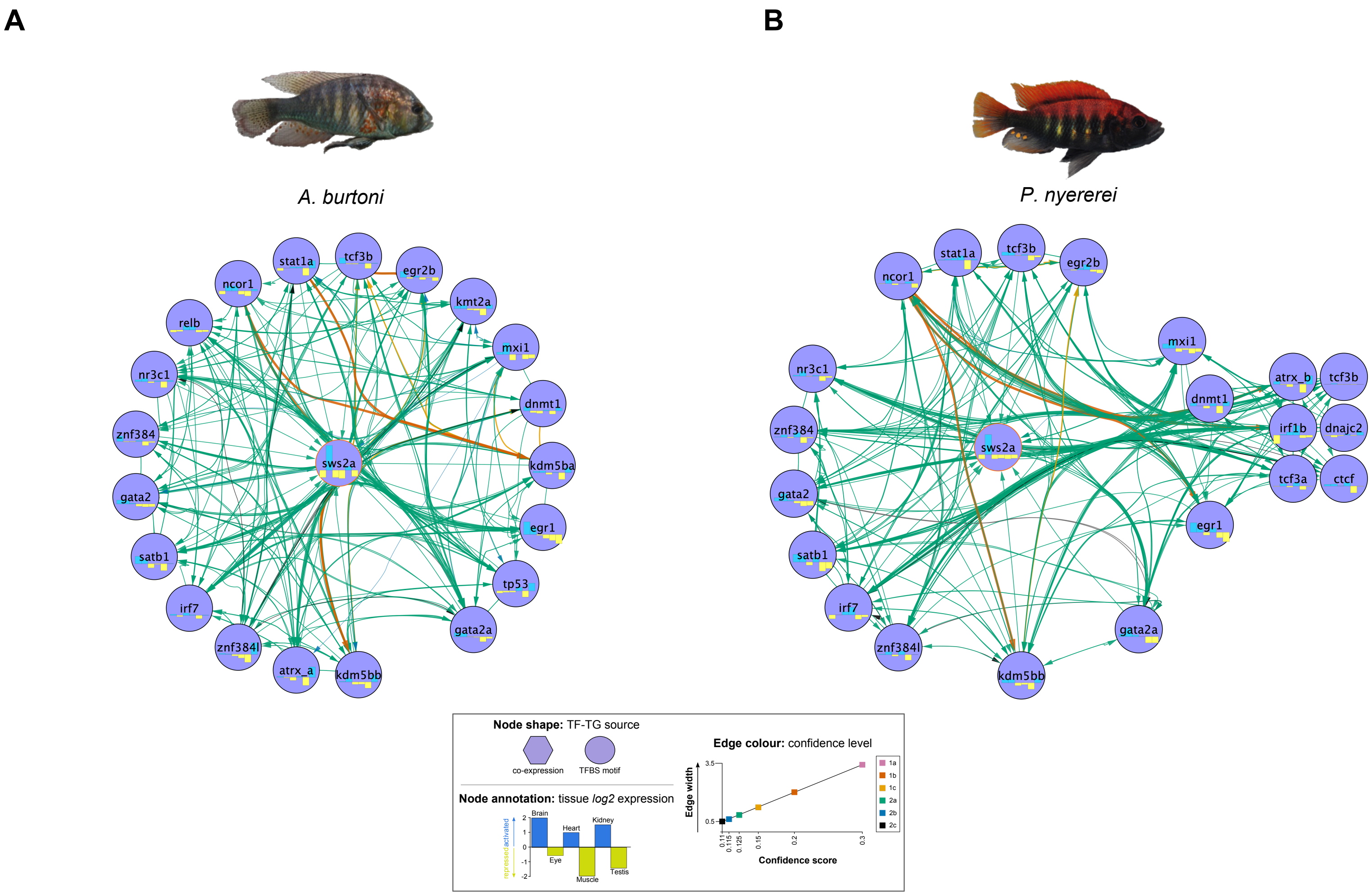


**Supplementary Figure S-R4a - Evolution of the *sws2a* opsin regulatory networks in *A. burtoni* and *P. nyererei.*** Regulatory networks of *sws2a* opsin shown for (A) *A. burtoni* and (B) *P. nyererei;* circular layout nodes are common regulators, grid layout nodes are unique regulators and node shape, annotation and edge color are denoted in legend.

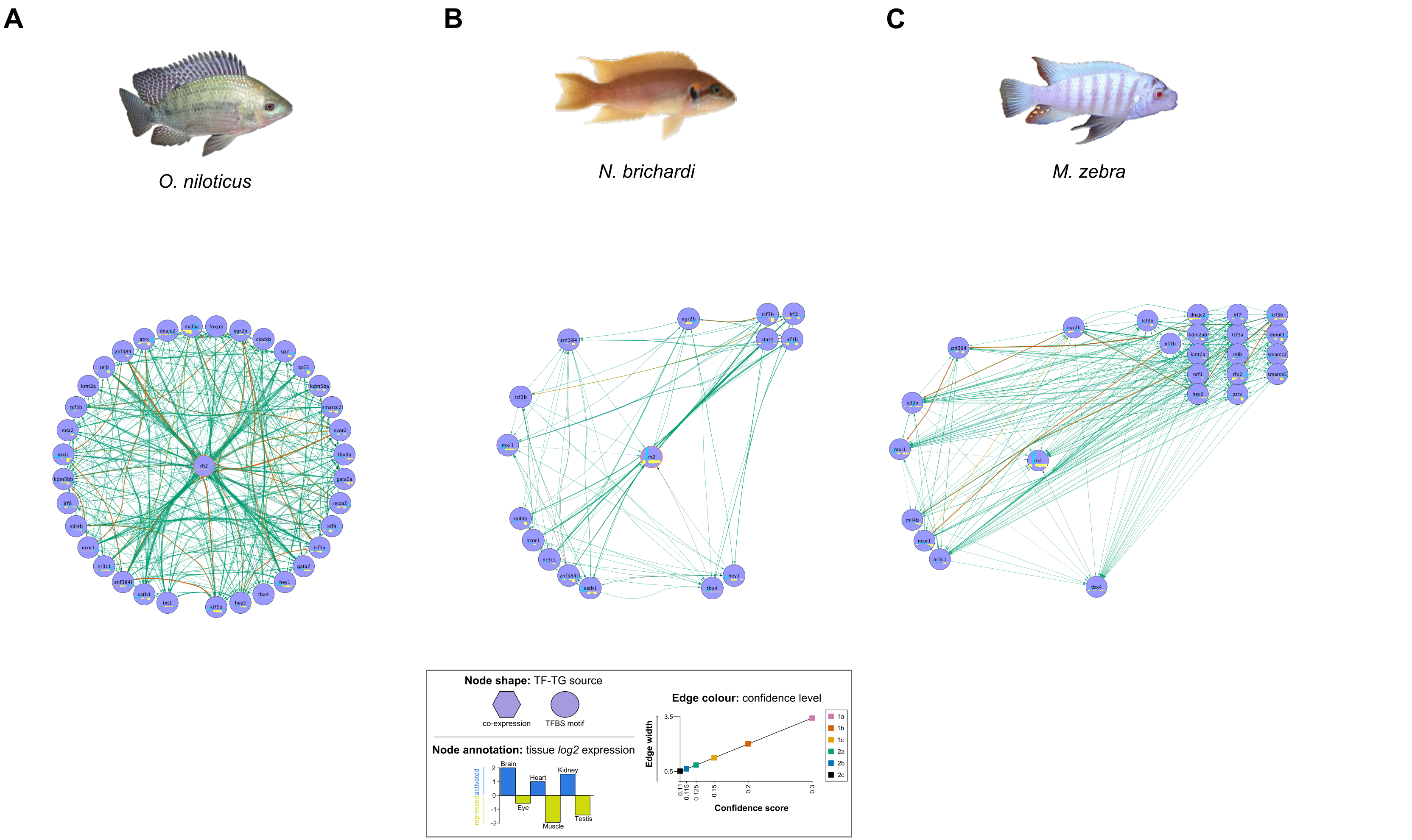


**Supplementary Figure S-R4b - Evolution of the *rh2b* opsin regulatory networks in *O. niloticus, N. brichardi* and *M. zebra*.** Regulatory networks of *rh2b* opsin shown for (A) *O. niloticus;* (B) *N. brichardi;* and (C) *M. zebra*; circular layout nodes are common regulators, grid layout nodes are unique regulators and node shape, annotation and edge color are denoted in legend.

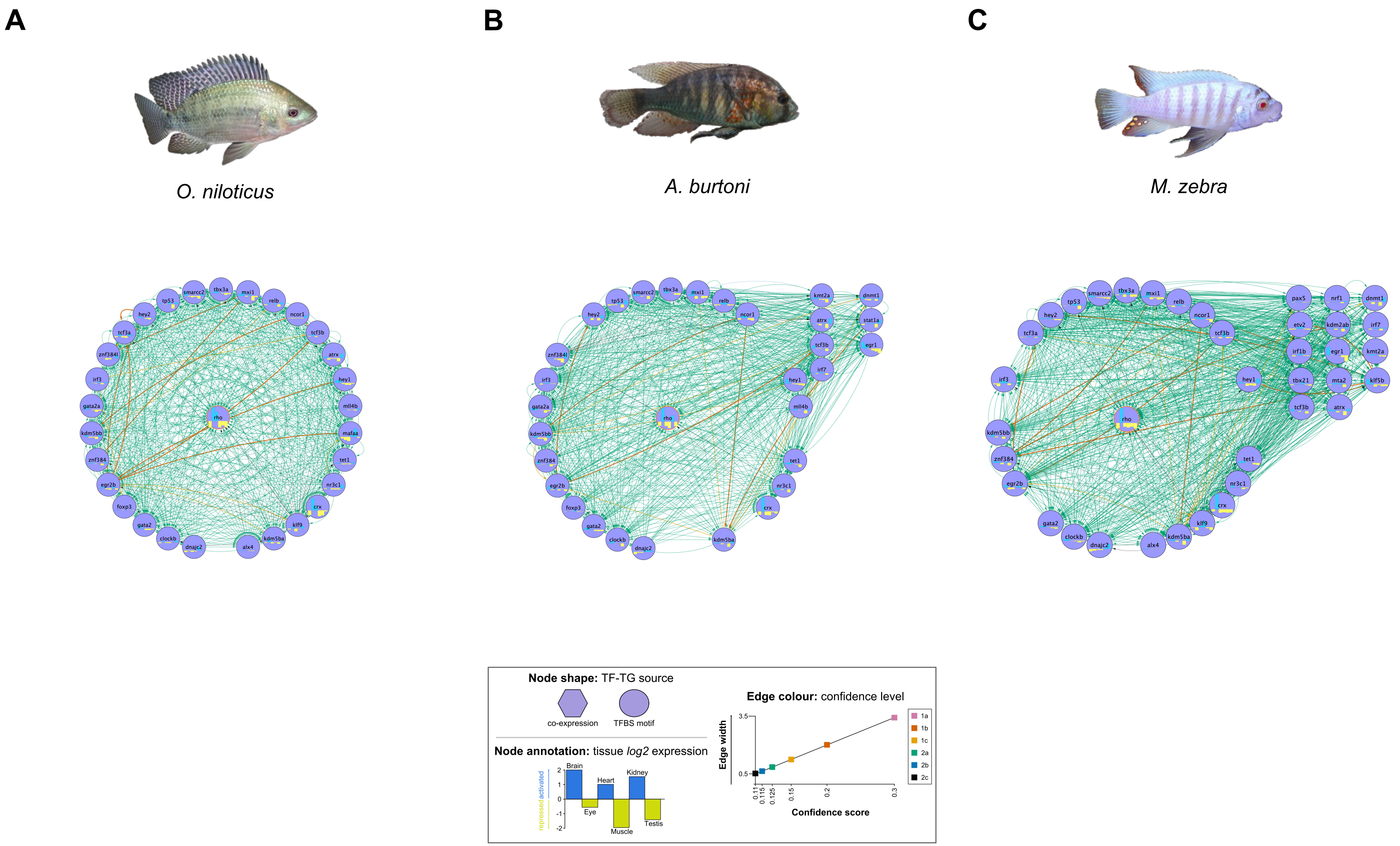


**Supplementary Figure S-R4c - Evolution of the rod opsin regulatory networks in *O. niloticus, A. burtoni* and *M. zebra.*** Regulatory networks of *rho* (rod) opsin shown for (A) *O. niloticus;* (B) *A. burtoni;* and (C) *M. zebra*; circular layout nodes are common regulators, grid layout nodes are unique regulators and node shape, annotation and edge color are denoted in legend.

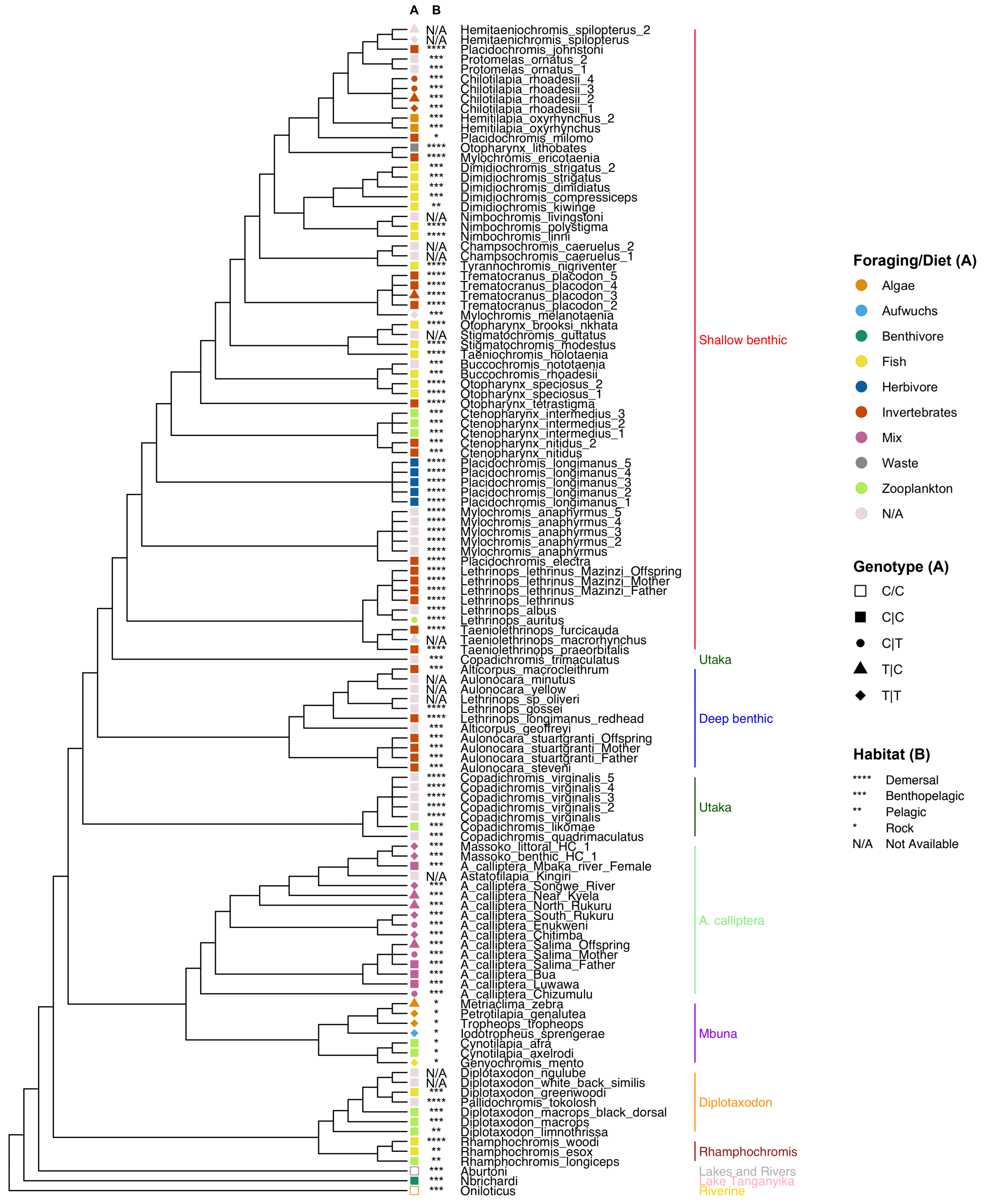


**Supplementary Figure S-R4d - Variants overlapping the Gata2a TFBS in *M. zebra rho* promoter, other Lake Malawi species and *O. niloticus, N. brichardi* and *A. burtoni* outgroups.** Lake Malawi phylogeny reproduced from published least controversial and all included species ASTRAL phylogeny[18]. Phylogenetic branches are labelled with species sample name and clade and according to legends (*right*): A) Species foraging/diet habit (color)[25] and phased SNP genotype (shape)[18]; B) species habitat[25,26].

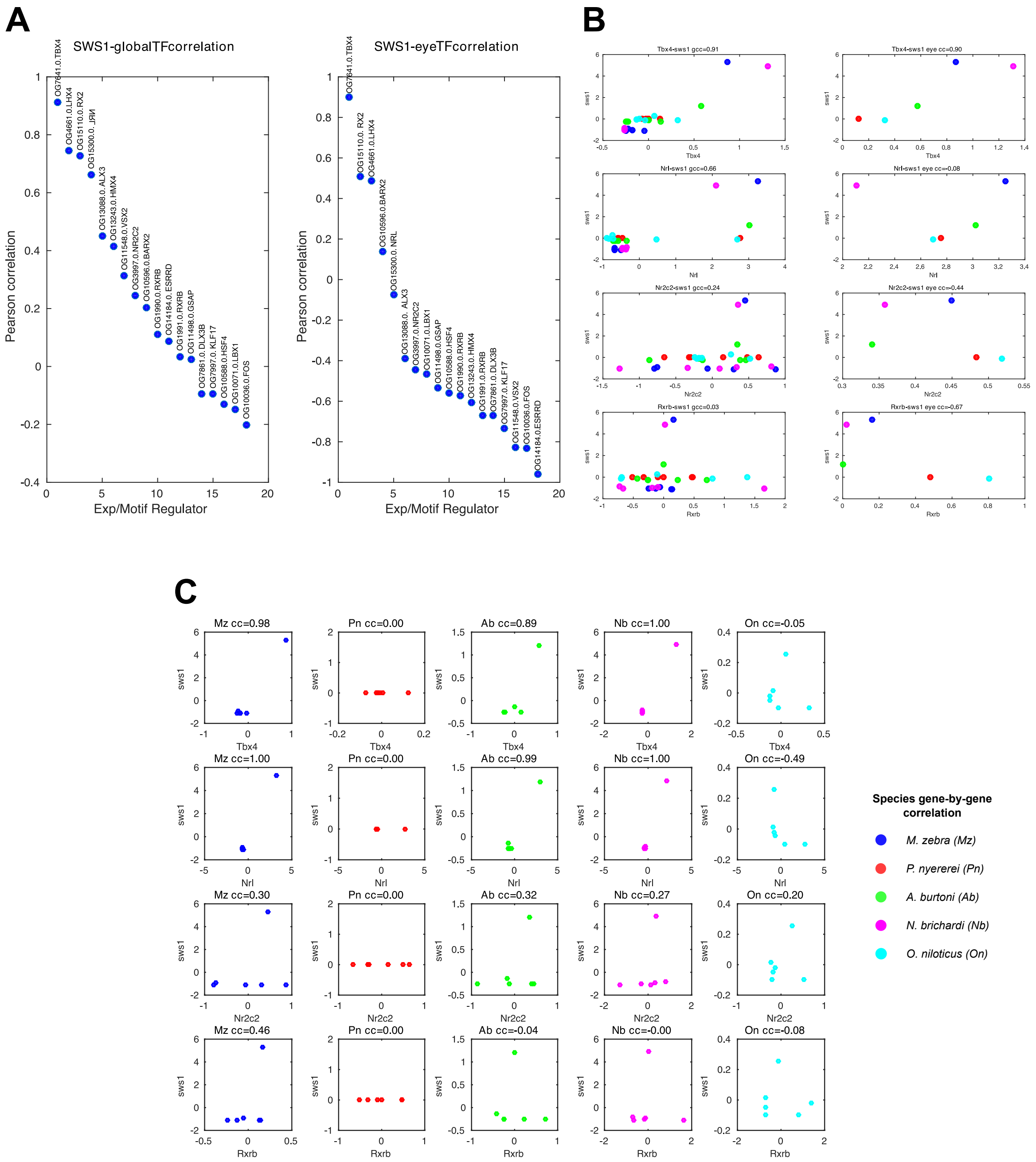


**Supplementary Figure S-R4e – Global and eye-specific correlations of *sws1* and primary interacting TFs expression. (A)** shows the Pearson correlation coefficient (y-axis) of all regulators expression separated by the expression of all species tissues (left) and eye tissue only (right). Scatter plots showing the expression level of the *sws1* gene (y-axis) and one of four possible regulators (x-axis), separated by **(B)** global tissue expression (left), eye expression (right) in all five species; and **(C)** expression in six tissues separated by each of the five species. Each dot corresponds to the expression of *sws1* and its regulator either in all tissues (Panel B, *left*; Panel C) or eye tissue only (Panel B, *right*). The Pearson's correlation coefficient (cc) between the regulator and *sws1*'s expression are mentioned in the title for panel B and C. For *sws1*, the first two sets of plots for TBX4 and NRL are among the top 2 regulators that were identified using our expression-based network inference approach. The remaining two rows, NR2C2 and RXRB were predicted based on motif scanning on the *sws1* gene promoter.

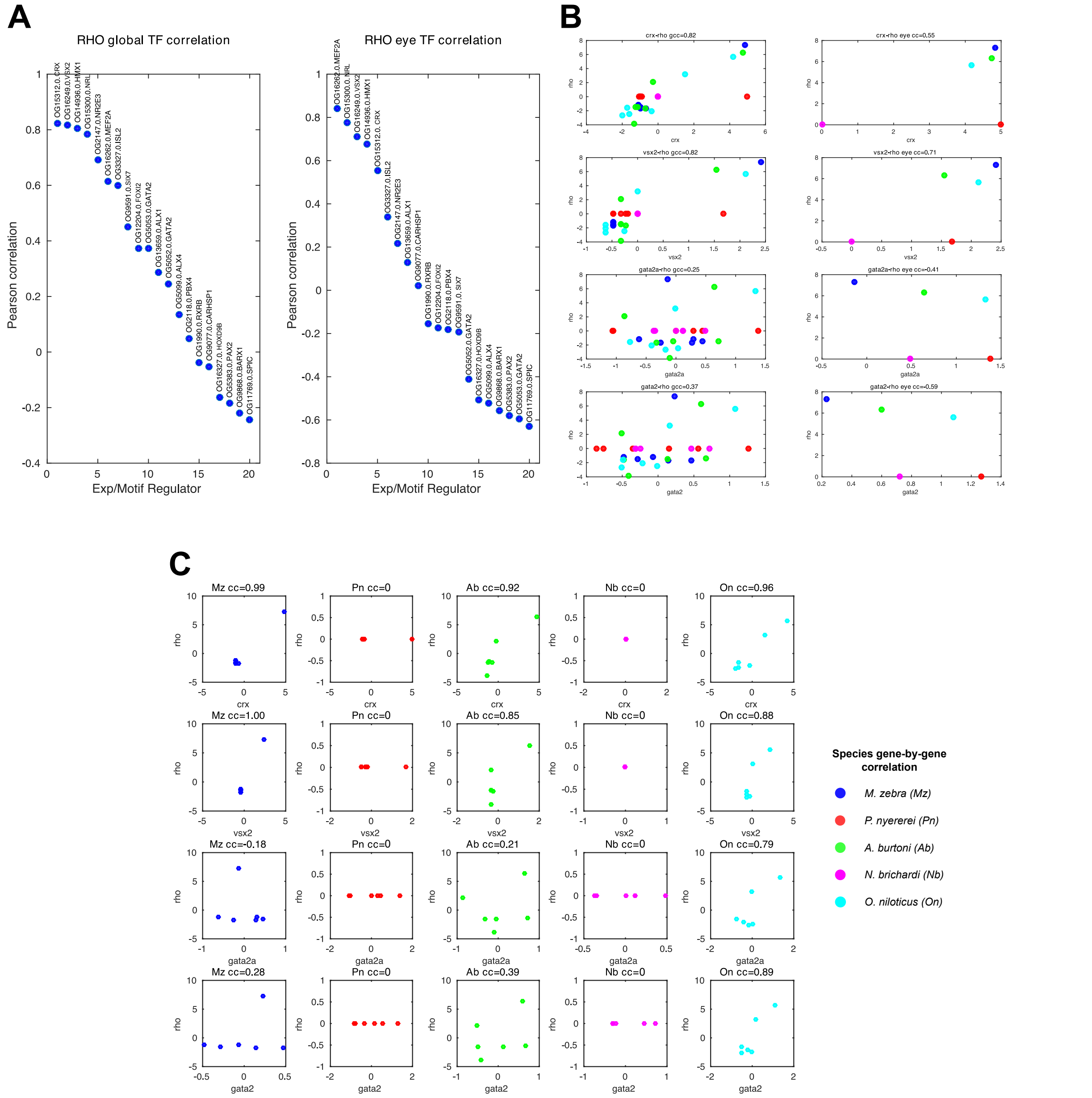


**Supplementary Figure S-R4f – Global and eye-specific correlations of *rho* and primary interacting TFs expression. (A)** shows the Pearson correlation coefficient (y-axis) of all regulators expression separated by the expression of all species tissues (left) and eye tissue only (right). Scatter plots showing the expression level of the *rho* gene (y-axis) and one of four possible regulators (x-axis), separated by **(B)** global tissue expression (left), eye expression (right) in all five species; and **(C)** expression in six tissues separated by each of the five species. Each dot corresponds to the expression of *rho* and its regulator either in all tissues (Panel B, *left*; Panel C) or eye tissue only (Panel B, *right*). The Pearson's correlation coefficient (cc) between the regulator and *rho*'s expression are mentioned in the title for panel B and C. For *rho*, the first two sets of plots for CRX and VSX2 are among the top 2 regulators that were identified using our expression-based network inference approach. The remaining two rows, GATA2A and GATA2 were predicted based on motif scanning on the *rho* gene promoter.

#### Methods

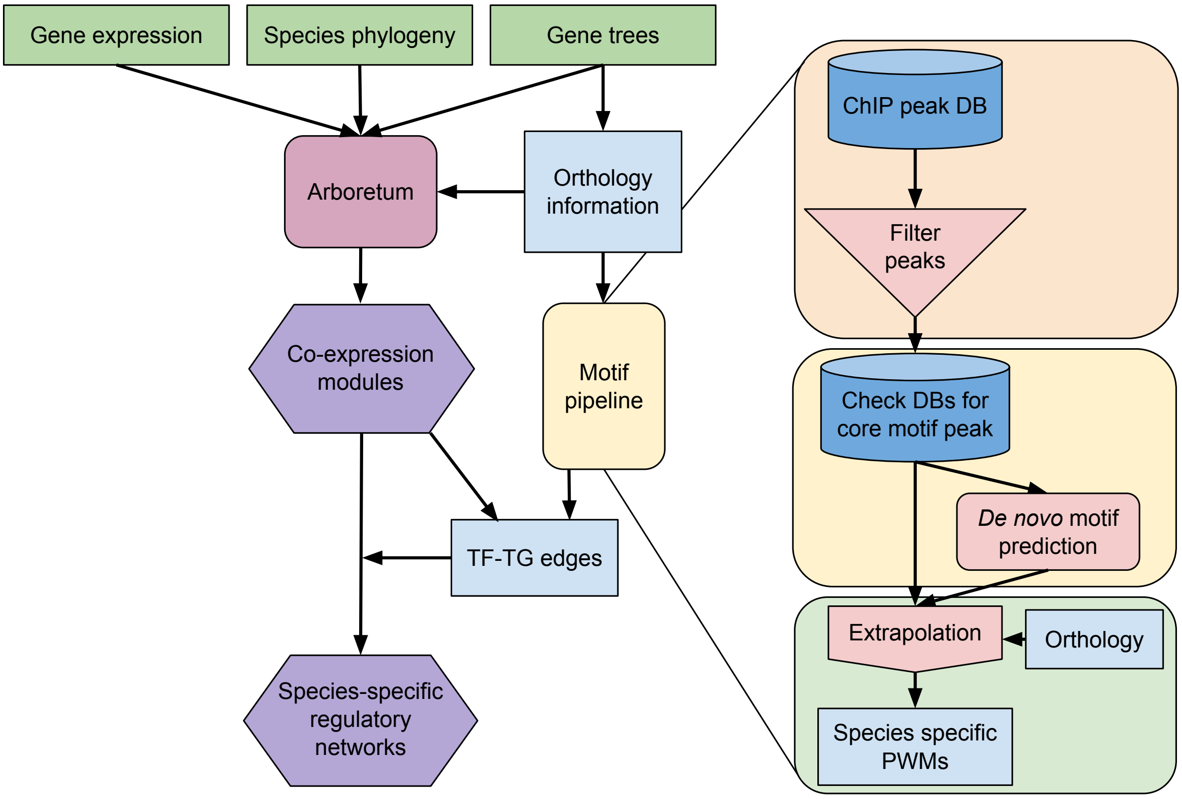


**Supplementary Figure S-M1 - Systematic framework for reconstructing and analyzing gene regulatory networks in five cichlids.** Our systematic framework comprises: (1) identifying modules of co-expressed genes from multi-tissue/multi-species and single-tissue/multi-species data using Arboretum[27]; (2) Using our developed motif prediction pipeline (*right)*, we integrate several datasets (co-expression, *cis* regulatory elements and transcription factor binding sites (TFBSs)) to; (3) reconstruct species-specific gene regulatory networks (GRNs) refined with gene expression data to find fine-grained tissue-specific network modules.

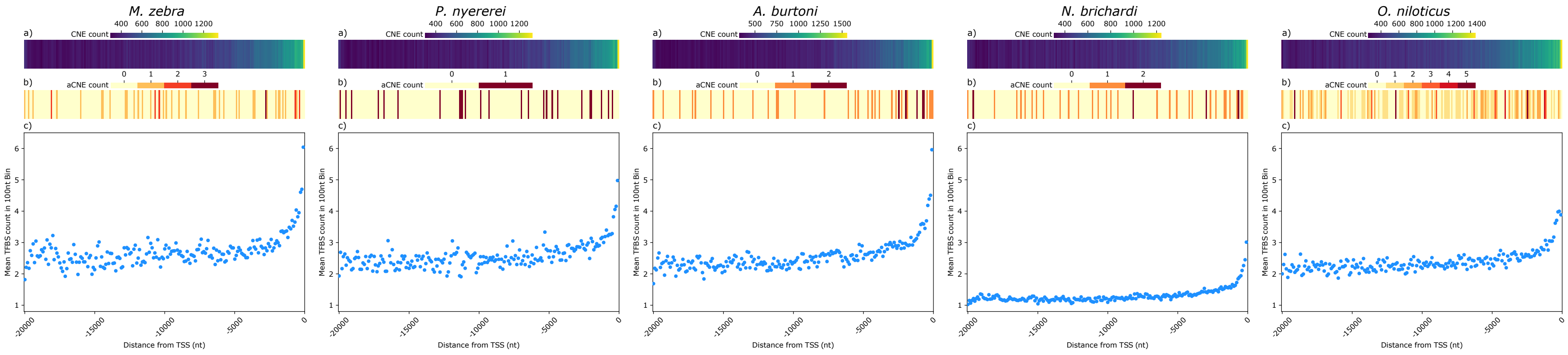


**Supplementary Figure S-M2. Counts of predicted TFBS in 100nt windows over 20kb regions upstream of the TSS in 5 cichlid species** shown below counts of conserved non-coding elements (CNEs) and CNEs that significantly diverged from the neutral model (aCNEs) over the same regions (see Brawand, D. *et al.* 2014 *Nature*). Heatmaps represent total counts of a) CNEs and b) aCNEs 0base midpoints intersecting 100nt windows for 20kb upstream of the TSS in each species. CNE and aCNE annotations used in a) and b) are taken from Brawand, D. *et al.* 2014 *Nature*, and were intersected with the 20kb *cis* regions described below before counting. Each data point in c) represents the total count of TFBS predictions within a 100nt window divided by the number of genes with a *cis* region contributing to that bin. *Cis* regions were called as described in the methods, briefly - if a gene was within 20kb to another gene annotation, the *cis* region called from the TSS was truncated to the boundary of the other annotation if on the same strand, or to an equal midpoint between annotations if on opposite strands, otherwise a full 20kb from the TSS was called. TFBSs were counted in a bin if the 0-base midpoint of the prediction fell within the bounds of the bin. To generate TFBS predictions, these 20kb *cis* regions were scanned with the total, nonredundant set of core extrapolations and FIMO *de novo* TFBS predictions for the species, generated as described in the *Methods*.

### Supplementary tables

| | | |
|---|---|---|
| **Nr2c2 DNA-binding Domain (DBD)** | | |
| *Primer* | *Sequence 5' > 3'* | *Notes* |
| mz-nb_T7_nr2c2DBD_F1 | GGATCCTAATACGACTCACTATAGGGAACAGCCACCATGTCAGGAGATTTGAGCCGACCA | Primer can be used for amplification in both Mz and Nb as it does not cover variable regions. There are only 4 (degenerate) mismatched bases between Nb and Mz; still coding for a 100% conserved DBD. |
| mz-nb_T7_nr2c2DBD_R1 | TTAGGGCACAATGTCGATGGGTTTCCTCTCACTCTGGACAGACTCAGTCTTCATCCCCAT | |
| **Rxrb DNA-binding Domain (DBD)** | | |
| mz-nb_T7_rxrbDBD_F1 | GGATCCTAATACGACTCACTATAGGGAACAGCCACCATGGCTCACAGCCCGGGAATAATG | Primer can be used for amplification in both Mz and Nb as it does not cover variable regions. There are only 3 (degenerate) mismatched bases between Nb and Mz; still coding for a 100% conserved DBD. |
| mz-nb_T7_rxrbDBD_R1 | TTACTGTCGTTCCTCTTGTACCGCTTCCCTCTTCATTCCCATGGCCAGGCACTTCTGGTA | |
| **DNA probes** | | |
| mz_cy5_nr2c2_rxrb-sws1TG_F1 | TCATTAGTCAGAGTCAGAGGTCACAGGA | Since the TFBS is predicted to be shared by both nr2c2 and rxrb, only a single DNA probe can be used to test binding of both nr2c2 and rxrb DBDs.<br><br>Negative control oligos are scrambled motifs of the original sequence so as to maintain nucleotide composition. |
| mz_nr2c2_rxrb-sws1TG_R1 | TCCTGTGACCTCTGACTCTGACTAATGA | |
| nb_cy5_nr2c2_rxrb-sws1TG_F1 | TCATTAGTCAGGGTCAGAGGTCACAGGA | |
| nb_nr2c2_rxrb-sws1TG_R1 | TCCTGTGACCTCTGACCCTGACTAATGA | |
| nb_cy5_nr2c2_rxrb-sws1TG-ve_F1 | TTGGTGAAAAGTGACTGGCTGCAGCAAC | |
| nb_nr2c2_rxrb-sws1TG-ve_R1 | GTTGCTGCAGCCAGTCACTTTTCACCAA | |

**Table S-M1a. Oligonucleotides used for amplification of DNA-binding domains (DBDs) and EMSA DNA probes.** The names and sequences are listed as pairs according to their usage.

| Reference Species_ Comparison Species | Total variants | Three Prime UTR | aCNEs | hCNEs | Non-State-changed Promoters | State-changed Promoters | TFBSs | Non-State-changed Promoters TFBSs | State-changed Promoters TFBSs | Promoter flank | Other variants |
|---|---|---|---|---|---|---|---|---|---|---|---|
| *mz11_pn1* | 7891416 | 197764 | 208 | 407996 | 610173 | 112568 | 1895509 | 1393708 | 501801 | 673391 | 6562707 |
| *mz11_ab1* | 9051743 | 230630 | 217 | 467713 | 696693 | 130742 | 2290382 | 1690816 | 599566 | 766937 | 7525748 |
| *mz11_nb1* | 18774705 | 537245 | 346 | 1026501 | 1449107 | 299816 | 3824346 | 2828340 | 996006 | 1622576 | 15461690 |
| *mz11_on11* | 32239921 | 1010838 | 368 | 1723742 | 2514873 | 550286 | 6062050 | 4449729 | 1612321 | 2853422 | 26439814 |
| *mz11_oryLat2* | 56596695 | 2239770 | 1175 | 7026932 | 3881344 | 866298 | 8379459 | 6164546 | 2214913 | 4923529 | 42581176 |
| *mz11_gasAcu1* | 64804903 | 3047571 | 1328 | 6872705 | 4506168 | 1082384 | 10632874 | 7735981 | 2896893 | 5736773 | 49294747 |
| *mz11_danRer7* | 24799703 | 622261 | 811 | 2802586 | 1487540 | 281167 | 1873134 | 1399465 | 473669 | 2155558 | 19605338 |
| *pn1_mz11* | 7891416 | 148832 | 652 | 524481 | 597081 | 118337 | 2076184 | 1512359 | 563825 | 632973 | 6502033 |
| *pn1_ab1* | 9474922 | 175499 | 628 | 584119 | 703023 | 143763 | 2306157 | 1671655 | 634502 | 751301 | 7867890 |
| *pn1_nb1* | 18837826 | 409594 | 761 | 1120740 | 1411056 | 312237 | 3803600 | 2767123 | 1036477 | 1556155 | 15583438 |
| *pn1_on11* | 31642942 | 754037 | 782 | 1773984 | 2409335 | 564952 | 5957661 | 4307448 | 1650213 | 2689308 | 26139852 |
| *pn1_oryLat2* | 55421413 | 1824348 | 1924 | 7072451 | 3733927 | 876813 | 8087791 | 5870042 | 2217749 | 4724492 | 41911950 |
| *pn1_gasAcu1* | 63474615 | 2442763 | 2323 | 6858563 | 4372824 | 1102629 | 10169486 | 7206188 | 2963298 | 5488028 | 48695513 |
| *pn1_danRer7* | 24243979 | 491588 | 1613 | 2915439 | 1475788 | 289715 | 1886208 | 1397056 | 489152 | 2083763 | 19069836 |
| *ab1_mz11* | 9051743 | 231059 | 550 | 617482 | 770949 | 138140 | 2543224 | 1856340 | 686884 | 788044 | 7293563 |
| *ab1_pn1* | 9474922 | 233267 | 550 | 615159 | 805441 | 144426 | 2413771 | 1779798 | 633973 | 817266 | 7676079 |
| *ab1_nb1* | 18755346 | 533312 | 667 | 1160524 | 1585165 | 308416 | 4046368 | 2957674 | 1088694 | 1668510 | 15167262 |
| *ab1_on11* | 31664404 | 989531 | 667 | 1837439 | 2692342 | 553697 | 6375972 | 4693714 | 1682258 | 2879081 | 25590728 |
| *ab1_oryLat2* | 55577322 | 2301485 | 1783 | 7315599 | 4152372 | 865748 | 8524348 | 6282664 | 2241684 | 5031746 | 40940335 |
| *ab1_gasAcu1* | 63706777 | 3057942 | 1975 | 7100149 | 4805932 | 1081607 | 10791519 | 7845950 | 2945569 | 5889798 | 47659172 |
| *ab1_danRer7* | 24308154 | 702771 | 1553 | 3052445 | 1575078 | 289508 | 2011002 | 1499144 | 511858 | 2210833 | 18686799 |
| *nb1_mz11* | 18774705 | 398795 | 854 | 1301826 | 1337025 | 317229 | 2787465 | 1901920 | 885545 | 1530873 | 15418976 |
| *nb1_pn1* | 18837826 | 399483 | 853 | 1287676 | 1339698 | 318890 | 2657360 | 1816406 | 840954 | 1538196 | 15491226 |
| *nb1_ab1* | 18755346 | 396301 | 837 | 1282981 | 1335049 | 318257 | 2670287 | 1809999 | 860288 | 1530025 | 15421921 |
| *nb1_on11* | 30695242 | 728828 | 813 | 1925303 | 2215016 | 559320 | 3997080 | 2714678 | 1282402 | 2575948 | 25265962 |

| | | | | | | | | | | | |
|---|---|---|---|---|---|---|---|---|---|---|---|
| *nb1_oryLat2* | 54190417 | 1793433 | 2187 | 7494458 | 3714716 | 884265 | 5205418 | 3545293 | 1660125 | 4629695 | 40301358 |
| *nb1_gasAcu1* | 62161124 | 2366297 | 2657 | 7315298 | 4316790 | 1095410 | 6590051 | 4443651 | 2146400 | 5396192 | 47064672 |
| *nb1_danRer7* | 23639358 | 519443 | 2019 | 3263109 | 1477914 | 293462 | 1244362 | 863477 | 380885 | 2068726 | 18083411 |
| *on11_mz11* | 32239921 | 1190969 | 530 | 1582304 | 2724962 | 518287 | 6569717 | 4810884 | 1758833 | 3069068 | 26222869 |
| *on11_pn1* | 31642942 | 1178735 | 520 | 1548053 | 2662480 | 511247 | 6196776 | 4534352 | 1662424 | 3011485 | 25741907 |
| *on11_ab1* | 31664404 | 1175171 | 531 | 1547076 | 2666456 | 511971 | 6197384 | 4568469 | 1628915 | 3012662 | 25763199 |
| *on11_nb1* | 30695242 | 1149444 | 496 | 1532045 | 2580351 | 499713 | 6036531 | 4430817 | 1605714 | 2924384 | 24933193 |
| *on11_oryLat2* | 55525414 | 2572618 | 1899 | 6592191 | 4263518 | 829077 | 9196930 | 6816539 | 2380391 | 5207882 | 41266111 |
| *on11_gasAcu1* | 63636292 | 3447429 | 2199 | 6469516 | 4991043 | 1064329 | 7589059 | 4443651 | 3145408 | 6129422 | 47661776 |
| *on11_danRer7* | 24264982 | 827485 | 1482 | 2520426 | 1560561 | 238361 | 1843646 | 1358928 | 484718 | 2182041 | 19116667 |

**Supplementary Table S-R2a - Species pairwise variants overlapping various genomic regions.** Pairwise variant calling and overlap to various genomic regions defined in *Methods.* Five cichlid species (mz11 – *M. zebra;* pn1 – *P. nyererei;* ab1 – *A. burtoni;* nb1 – *N. brichardi;* and on11 – *O. niloticus*) and three outgroup teleost species (oryLat2 - medaka, gasAcu1 - stickleback and danRer7 - zebrafish) defined within.

| Species | No. of gene nodes | Co-expressed TF-TG edges | TF-TG (promoter TFBS) edges |
|---|---|---|---|
| *M. zebra* | 11,075 | 3,964 | 4,760,610 |
| *P. nyererei* | 11,070 | 4,029 | 4,862,871 |
| *A. burtoni* | 11,638 | 3,822 | 5,355,927 |
| *N. brichardi* | 10,015 | 3,180 | 3,292,032 |
| *O. niloticus* | 11,790 | 4,099 | 5,896,075 |

**Supplementary Table S-R3a - Number of statistically significant edges in species reconstructed networks**. Network edges derived from several sources (see *Methods*), representing various regulatory interactions/associations in the five cichlids.

#### Extended data legends

**Extended Data S-R1A** - **Gene Ontology (GO) enrichment of modules across all extant and ancestral species.** Enriched terms of significance FDR-corrected *P*-value (*q*-value <0.05) in modules (rows and ':n' module number) are shown for extant and ancestral species (columns) and colored according to module and gradient, -log(*q*-value) in each grid position (see legend, *left*). Set-based hypergeometric test of enrichment carried out using a background of all module genes.

**Extended Data S-R1B** - **Transcription factor motif enrichment of module gene promoters across all extant and ancestral species.** All enriched motifs of significance FDR-corrected *P*-value (*q*-value <0.05) in modules (rows and ':n' module number) are shown for extant and ancestral species (columns) and colored according to module and gradient, -log(*q*-value) in each grid position (see legend, *left*). Set-based hypergeometric test of enrichment carried out using a background of all motifs predicted within all module gene promoters.

**Extended Data S-R1C-H** - **Heatmap matrices of enrichment, expression and Pearson correlation between the two for each motif, across all six tissues (brain, eye, heart, kidney, muscle and testis).** First five columns are gradient coloured (legend to *right*) according to enriched motifs of significance FDR-corrected *P*-value (*q*-value <0.05) in module gene promoters (rows as '_n' module number and '_TF' motifs) shown as *-log*(q-value) in all five extant species (columns). Next five columns are tissue-specific zero-mean log-expression expression ratios (legend to *right)* used as input for Arboretum (see *Methods*) in all five extant species (columns). Final column is Pearson correlation coefficient (PCC) of motif enrichment and expression values shown in previous columns. PCC ranges (legend to *right)* from 1 (positive linear correlation, blue), 0 (no linear correlation, white), and −1 (negative linear correlation, red).

**Extended Data Table S-R3A** - DyNet[12] rewiring scores of TF-TG 1-to-1 edges when all five species networks are compared.

**Extended Data Table S-R3B** - List of candidate genes from previous molecular evolutionary and developmental studies[2,13,14,25] potentially associated with phenotypic novelty in cichlids that overlap the six tissues we studied.

**Extended Data Table S-R3C** - DyNet[12] rewiring scores of candidate genes in TF-TG 1-to-1 edges when all five species networks are compared.

**Extended Data Table S-R3D** - DyNet[12] rewiring scores of TF-TG all edges when all five species networks are compared.

**Extended Data Table S-R3E** - DyNet[12] rewiring scores of candidate genes in TF-TG all edges when all five species networks are compared.

**Extended Data Table S-R3F** – Rate of edge gain and loss in TF-TG all edges across the five cichlid phylogeny. Rates that are >100 (and excluded from analyses) are rank shaded blue (gain) and green (loss) whereas likelihood scores are rank shaded orange.

12. Goenawan IH, Bryan K, Lynn DJ. DyNet: Visualization and analysis of dynamic

molecular interaction networks. Bioinformatics. 2016. p. 2713–5.
13. Brawand D, Wagner CE, Li YI, Malinsky M, Keller I, Fan S, et al. The genomic substrate for adaptive radiation in African cichlid fish. Nature. 2014;93:17–9.
14. Hahn C, Genner MJ, Turner GF, Joyce DA. The genomic basis of cichlid fish adaptation within the deepwater "twilight zone" of Lake Malawi. Evol Lett. 2017;1:184–98.
15. Whited JL. Dynactin is required to maintain nuclear position within postmitotic Drosophila photoreceptor neurons. Development. 2004;131:4677–86.
16. Cohen DR, Cheng CW, Cheng SH, Hui CC. Expression of two novel mouse Iroquois homeobox genes during neurogenesis. Mech Dev. 2000;91:317–21.
17. Choy SW, Cheng CW, Lee ST, Li VWT, Hui MNY, Hui CC, et al. A cascade of irx1a and irx2a controls shh expression during retinogenesis. Dev Dyn. 2010;239:3204–14.
18. Malinsky M, Svardal H, Tyers AM, Miska EA, Genner MJ, Turner GF, et al. Whole-genome sequences of Malawi cichlids reveal multiple radiations interconnected by gene flow. Nat Ecol Evol. 2018;2:1940–55.
19. Conte MA, Kocher TD. An improved genome reference for the African cichlid, Metriaclima zebra. BMC Genomics. 2015;16.
20. Conte MA, Joshi R, Moore EC, Nandamuri SP, Gammerdinger WJ, Roberts RB, et al. Chromosome-scale assemblies reveal the structural evolution of African cichlid genomes. Gigascience. 2019;
21. Musser JM, Arendt D. Loss and gain of cone types in vertebrate ciliary photoreceptor evolution. Dev. Biol. 2017.
22. Mears AJ, Kondo M, Swain PK, Takada Y, Bush RA, Saunders TL, et al. Nrl is required for rod photoreceptor development. Nat Genet. 2001;
23. Evans RM, Mangelsdorf DJ. Nuclear receptors, RXR, and the big bang. Cell. 2014. p. 255–66.
24. Olivares AM, Moreno-Ramos OA, Haider NB. Role of nuclear receptors in central nervous system development and associated diseases. J. Exp. Neurosci. 2015.
25. Hofmann CM, O'Quin KE, Justin Marshall N, Cronin TW, Seehausen O, Carleton KL.

The eyes have it: Regulatory and structural changes both underlie cichlid visual pigment diversity. PLoS Biol. 2009;7.

26. Froese R, Pauly D. Fishbase [Internet]. FishBase. 2017. Available from: www.fishbase.org

27. Roy S, Wapinski I, Pfiffner J, French C, Socha A, Konieczka J, et al. Arboretum: Reconstruction and analysis of the evolutionary history of condition-specific transcriptional modules. Genome Res. 2013;23:1039–50.
